# Machine learning-based prediction of fish acute mortality: Implementation, interpretation, and regulatory relevance

**DOI:** 10.1101/2024.03.14.584983

**Authors:** Lilian Gasser, Christoph Schür, Fernando Perez-Cruz, Kristin Schirmer, Marco Baity-Jesi

**Affiliations:** Swiss Data Science Center (SDSC), Andreasstrasse 5, Zürich, Switzerland; Eawag, Swiss Federal Institute of Aquatic Science and Technology, Dübendorf, Switzerland; ETH Zürich: Department of Computer Science, Zürich, Switzerland; ETH Zürich: Department of Environmental Systems Science, Zürich, Switzerland

## Abstract

Regulation of chemicals requires knowledge of their toxicological effects on a large number of species, which has traditionally been acquired through *in vivo* testing. The recent effort to find alternatives based on machine learning, however, has not focused on guaranteeing transparency, comparability and reproducibility, which makes it difficult to assess advantages and disadvantages of these methods. Also, comparable baseline performances are needed. In this study, we trained regression models on the ADORE “t-F2F” challenge proposed in [Schür *et al*., *Nature Scientific data*, 2023] to predict acute mortality, measured as LC50 (lethal concentration 50), of organic compounds on fishes. We trained LASSO, random forest (RF), XGBoost, Gaussian process (GP) regression models, and found a series of aspects that are stable across models: (i) using mass or molar concentrations does not affect performances; (ii) the performances are only weakly dependent on the molecular representations of the chemicals, but (iii) strongly on how the data is split. Overall, the tree-based models RF and XGBoost performed best and we were able to predict the log10-transformed LC50 with a root mean square error of 0.90, which corresponds to an order of magnitude on the original LC50 scale. On a local level, on the other hand, the models are not able to consistently predict the toxicity of individual chemicals accurately enough. Predictions for single chemicals are mostly influenced by a few chemical properties while taxonomic traits are not captured sufficiently by the models. We discuss technical and conceptual improvements for these challenges to enhance the suitability of *in silico* methods to environmental hazard assessment. Accordingly, this work showcases state-of-the-art models and contributes to the ongoing discussion on regulatory integration.

**Environmental significance:** Conventional environmental hazard assessment in its current form will not be able to adapt to the growing need for toxicity testing. Alternative methods, such as toxicity prediction through machine learning, could fulfill that need in an economically and ethically sound manner. Proper implementation, documentation, and the integration into the regulatory process are prerequisites for the usability and acceptance of these models.

## 1 Introduction

Chemical regulation aims to ensure the safety of humans and the environment, which is traditionally based on animal testing. As an example, in the European Union, the legislation for the Registration, Evaluation, Authorisation and Restriction of Chemicals (REACH) ^1^ requires (invertebrate) animal tests to be performed for chemicals with a yearly import or production volume of more than 1 ton. Acute (*i*.*e*., short-term) fish mortality tests are required for chemicals with an import or production volume of 10 tons *per annum* or more and are standardized through the OECD test guideline (TG) 203^2^. The global use of birds and fish was estimated to range between 440,000 and 2.2 million individuals at a cost upwards of $39 million *per annum* ^3^. Consequently, reducing the use of animals and, more specifically, fish acute toxicity testing has a high priority in chemical hazard assessment, both from an economical and ethical perspective.

In the past decade, there has been an increased effort towards the adoption of new approach methods (NAMs), *i*.*e*., implementing and validating alternative methods to move away from measuring chemical toxicity *in vivo* with the sacrifice of animals. Computer-based (*in silico*) methods have the potential to supplement, if not replace, animal testing through predictive toxicology based on historical data ^4^. Increased computational power, accessibility and ease of use of software, and recognition of the potential by legislators has contributed to increased research efforts of *in silico* alternative methods.

Quantitative structure activity relationship (QSAR) modeling is the umbrella term for models based on the similarity-property principle, *i*.*e*., the assumption that chemicals with similar structure will elicit a similar biological response. In the field of toxicity, these models are sometimes referred to as quantitative structure toxicity relationship (QSTR) models, which have a long history of predicting toxicological outcomes using either linear or nonlinear relationships between chemical descriptors and a biological response ^5^. More than 10,000 QSAR models were published or publicly available in 2023^6^. Recently, QSAR research has started to incorporate machine learning (ML), *i*.*e*., computational methods that are able to find hidden patterns in large amounts of data without explicit programming and, on the basis of said patterns, are able to make predictions. The application of ML comes with the caveat that domain-experts are not necessarily also ML experts.

QSARs are characterized by the relationship they are applied to, *i*.*e*., the chemical descriptor(s) used to predict a biological outcome, and not by the underlying modeling approach. Hence, integrating information beyond chemical descriptors is not adequately captured by the term QSAR. Zubrod *et al*. (2023) referred to models also including species-specific and experimental information as Bio-QSARs ^7^. ML methods can be applied to both QSARs and extended QSARs with non-chemical features.

So far, mammal-centered toxicology was the primary focus of ML-based predictive toxicology. Notably, Luechtefeld *et al*. (2018) implemented read-across structure activity relationship (RASAR) based on binary fingerprints and Jaccard distances, which they applied to different endpoint groups ^8^. They compared the model performance to the variability of the *in vivo* data, which they found to be similar, although their inclusion of modeled data and lack of transparent reporting and data availability have been criticized ^9^. In their response to the critique, the authors explicitly point out that their approach differs from QSAR by the use of big data and artificial intelligence as opposed to small and highly curated datasets and conclude that certain criticisms to QSARs do not apply to their approach ^10^. Wu *et al*. (2022) brought Luechtefeld *et al*.’s approach to the realm of ecotoxicology by applying it to toxicity classification of acute fish mortality ^11^ and found that their RASAR models did not outperform random forest (RF) models.

Despite getting less attention of ML than mammal-centered toxicology, several studies predict ecotoxicological outcomes using regression. They differ in the employed approaches, most notably in the datasets used and, therefore, in the chemical and taxonomic spaces ^7,12–14^.

Nevertheless, the adoption of ML in ecotoxicological research is still in its infancy, which comes with inherent pitfalls. Data leakage, one of the most common issues when applying ML models, “is a spurious relationship between the independent variables and the target variable that arises as an artifact of the data collection, sampling, or pre-processing strategy. Since the spurious relationship won’t be present in the distribution about which scientific claims are made, leakage usually leads to inflated estimates of model performance.” ^15^ It arises when data points from repeated measurements are assigned to both the training and the test set and results in the model merely recalling the relationship between the response and feature combinations instead of making a prediction based on a learned pattern. Data leakage can also occur when information about the response is introduced that should not legitimately be used for modeling ^16^. As of 2023, the issue of data leakage has been described to affect 329 papers across 17 research fields ^15^. Stock *et al*. (2023) discussed domain-specific risks of data leakage for the use of ML models in ecology and argued for the creation of domain-specific guidelines to avoid data leakage and related phenomena, such as short-cut learning ^17^.

Besides the issue of data leakage, predictive ecotoxicology lacks commonly recognized best practices such as benchmark datasets and reporting standards ^15,18^. As a part of ML-based research, it faces a reproducibility crisis, partly caused by inconsistent and in-transparent reporting (including underlying computer code), which prevents peer-reviewers from adequately assessing the findings, the modeling, and the data those findings are based on. Several efforts aim to sensitize researchers to common pitfalls ^20,21^ and to motivate them to adopt checklist-based reporting standards, such as REFORMS proposed by Kapoor *et al*. (2023) ^22^. For QSAR models, similar quality standards have already been proposed (with 49 assessment criteria covering various aspects of QSAR development, documentation and use) ^18^ and further developed specifically for the application of ML methods to QSARs ^19^. Furthermore, the FAIR (Findable, Accessible, Interoperable, Reusable) principles, which were developed for data sharing, could be adapted to model description and deployment and therefore help to improve the reproducibility and largescale adoption of these methods, and eventually turn them into a (re)usable resource for chemical safety assessment ^6^.

Data handling, *i*.*e*., curation, processing, and use in a modeling framework, plays an equally crucial role to avoid reproducibility issues. It requires both domain and ML expertise. Model applicability and performance highly depends on the data it was trained on. There is a trade-off between restrictive data filtering leading to narrowly applicable models, that are thus not very relevant, and unrestrictive data filtering yielding models that might cover a large range of species and chemicals, but are not accurate enough ^23^.

This paper is based on the well-characterized benchmark dataset ADORE for acute mortality in ecotoxicology. We investigate the application of ML methods to fish acute toxicity, namely the prediction challenge “t-F2F” on the taxonomic group fish covering 140 species and 1,905 chemicals. Six molecular representations are available to computationally represent molecules: the fingerprints MACCS, PubChem, Morgan, ToxPrint, the molecular descriptor Mordred, and the mol2vec embedding. Using this dataset allows to produce reproducible and comparable results that can act as a benchmark for future studies. We apply the four models LASSO, random forest, XGBoost, and Gaussian process regression. We train all combinations of molecular representations and models. We then analyse the model results to gain a better understanding of relevant features and aspects of the dataset. We aim to present state-of-the-art methods in an accessible manner for modeling experts, (eco)toxicologists, and regulators, alike. For the sake of transparency, we perform a self-assessment of the dataset, models, and reporting in accordance with proposed best practices ^15,19,22^.

## 2 Data

In this section, we introduce the data focusing on the relevant challenge, response values, features, and data splits.

### 2.1 Data generation and description

The benchmark dataset ADORE on acute mortality contains toxicity tests of three relevant taxonomic groups (fish, crustaceans, and algae) ^24^. The core of ADORE originates from the ECOTOX database ^25^, which was harmonized and pre-filtered to only contain entries suitable to model acute toxicity in fish, crustaceans, and algae. This core dataset was expanded with taxonomic and chemical information from various sources and then filtered to only contain entries on acute mortality for which information from all sources is available. The filtered dataset mainly contains entries from organic chemicals. In total, the ADORE dataset contains 33,448 entries, of which more than 75%, *i*.*e*., 26,114 entries are on fish, 6,630 entries on crustaceans, and 704 entries on algae. Please refer to the corresponding paper for a detailed description of the dataset ^24^. Here, we summarize the aspects relevant for this study.

### 2.2 Focus on fish challenge

The ADORE challenges on acute mortality cover three levels of complexity. The most complex challenges are based on the whole dataset including all three taxonomic groups (fish, crustaceans, and algae). At an intermediate level of complexity, challenges focus on one taxonomic group. Finally, the least complex challenges are restricted to single, well-represented test species. In this study, we focused on the taxonomic group of fish. Using the “t-F2F” challenge, we aimed to find the best combination of model and molecular representation with the corresponding features to predict acute mortality across 140 fish species.

### 2.3 Response values

The dataset contains only entries with the endpoint lethal concentration 50 (LC50) for fish mortality. All LC50 values were converted to *mg/L* and *mol/L*, and then log10-transformed. In this work, we predict both log molar and log mass LC50 (Fig. 1).

**Fig. 1.**
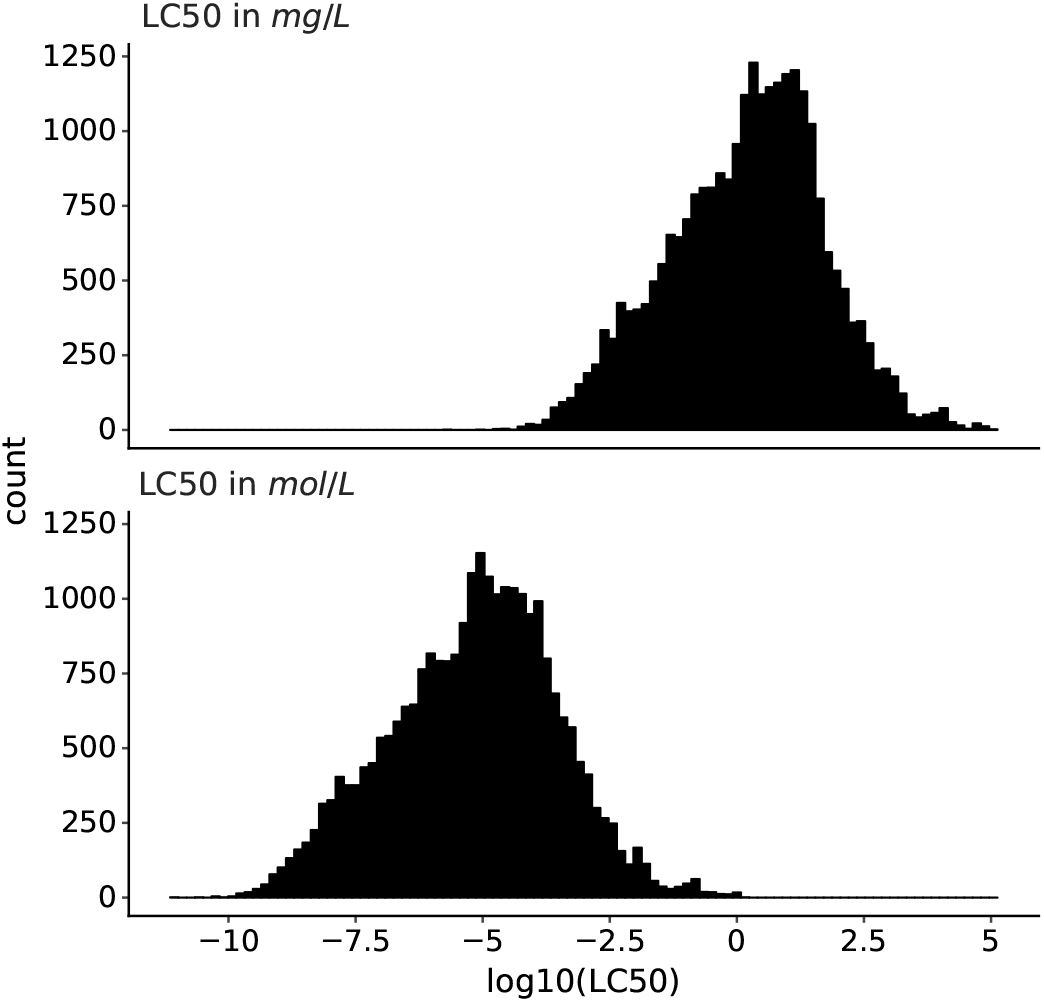
Histograms of log10-transformed LC50 (lethal concentration 50) in mass and molar units.

### 2.4 Description and processing of modeling features

The features can be summarized in three categories: experimental, chemical, and taxonomic. The responses and the modeling features are listed in the supplemental Table 1.

The *experimental features* describe the experimental conditions, specifically, observation duration, media type, exposure type, and concentration type that we used as the four experimental features in the models. The observation duration is ordinal with four levels (24, 48, 72, 96 hours), which were transformed to be in the range of [0, 1]. The other three features are categorical and were one-hot-encoded (*i*.*e*., translated to a binary vector for each level indicating its presence or absence. We do not deem the other experimental information included in the dataset relevant,, and for some features, we argue in Ref. ^24^ against using them for modeling.

The *chemical features* can be split in two sub-categories. Firstly, we include computable properties such as molecular weight (in *g/mol*), water solubility (in *mg/L*), melting point (in °*C*), and the octanol-water partition coefficient (*logK*_*ow*_, logP), for which positive/higher values indicate higher lipophilicity of a compound. We used these four features, standardized based on the training data, and opted against using the other computable features in the dataset, which are based on numbers of atoms, bonds and molecular substructures, as they are correlated with the selected features.

Secondly, the ADORE dataset contains six molecular representations, which were developed to make chemical structures machine-readable and therefore usable for ML models. The four fingerprints MACCS, PubChem, Morgan, and ToxPrint, as well as the molecular descriptor Mordred are examples of non-learned representations whereas mol2vec is a learned molecular embedding ^26^. Please refer to Schür *et al*. (2023) ^24^ for a detailed description. For including a fingerprint as model features, we suggest to remove duplicated and uninformative bits, (*i*.*e*., those with only little variation, following the modeling pipeline described in Lovric *et al*. ^27^). For the “t-F2F” dataset, the number of informative bits for the four fingerprints are shown in Table 1, determined for a standard deviation threshold of 0.1. The number of informative features are determined based only on the training data to avoid data leakage, which explains the different numbers for the two data splitting schemes. For mol2vec, we retained all 300 features, standardized based on the training data. For Mordred, we standardized the continuous features based on the training data and performed a uniform transformation of the ordinal features to the range of [0, 1].

**Table 1.** Feature count for each fingerprint and data splitting combination. Most features are from the molecular representations, see column *n*_*mol*.*repr*._, where we count the number of informative bits for the four fingerprints, give the dimensionality of the embedding for mol2vec, and the number of selected features for Mordred. The 37 remaining features are experimental, taxonomic, and chemical properties (*n*_*other*_). For Mordred, we do not use the four chemical properties as they are already part of the molecular descriptor.

| mol. repr. | data split | $n_{all}$ | $n_{mol.repr.}$ | $n_{other}$ |
| --- | --- | --- | --- | --- |
| MACCS | totally random | 180 | 143 | 37 |
| MACCS | occurrence | 178 | 141 | 37 |
| PubChem | totally random | 506 | 469 | 37 |
| PubChem | occurrence | 508 | 471 | 37 |
| Morgan | totally random | 422 | 385 | 37 |
| Morgan | occurrence | 417 | 380 | 37 |
| ToxPrint | totally random | 211 | 174 | 37 |
| ToxPrint | occurrence | 209 | 172 | 37 |
| mol2vec | totally random | 337 | 300 | 37 |
| mol2vec | occurrence | 337 | 300 | 37 |
| Mordred | totally random | 577 | 544 | 33 |
| Mordred | occurrence | 577 | 544 | 33 |

The *taxonomic features* can also be split in two sub-categories. Firstly, the Add my Pet database ^28^ provides features on ecology, life-history, and pseudo-data used for dynamic energy budget (DEB) modeling ^29^. From ecology, we included the categorical variables climate zone, ecozone, food, and the binary coded migratory behavior. For these categorical variables, we used a many-hot encoding since a fish may fulfill more than one level, *e*.*g*., for the fathead minnow, the entry for the food variable, “D_H”, means that it is both detrivorous (“D”) and herbivorous (“H”). From life-history, we included life span and ultimate body length. From pseudo-data, we included energy conductance, allocation rate to soma and volume-specific somatic maintenance cost. The continuous variables were standardized based on the training data.

Secondly, the ADORE dataset includes the phylogenetic distance between species to account for the genetic relationship between species that might be exploited to infer toxicity across species. This is based on the assumption that closely-related species will have a more similar sensitivity profile than less-closely related ones ^30^. The phylogenetic distance cannot be readily used in a standard model as it is a pairwise distance that cannot solely be attributed to a data point. We only used it in conjunction with GP regression.

Mainly, the models are trained on all these features except for the phylogenetic distances. We also trained the models without a molecular representation, *i*.*e*., using only experimental features, chemical, and taxonomic properties (abbreviated as ‘none’), and with only three chemical properties, namely molecular weight, water solubility, and logP (‘top 3’). Additionally, we trained GP models with all features including the phylogenetic distances.

### 2.5 Data splittings

Data splitting, the generation of training and test data subsets, and of cross-validations folds, can greatly affect model performance. Possible causes are the inherent variability of the data itself and (non-obvious) data leakage. Schür *et al*. (2023) discusses different data splitting schemes in detail ^24^. Here, we describe the two data splittings considered in this study.

#### 2.5.1 Split totally at random

The simplest train-test-split can be achieved by random sampling of data points, which has been the main approach in previous work applying ML to ecotoxicology and generally suffices for a well-balanced dataset without repeated experiments ^12,13, 31^. For the ADORE dataset with repeated experiments, *i*.*e*., data points coinciding in chemical, species, and experimental conditions, this approach has a high risk of data leakage and the associated overestimated model performances, as the same chemical as well as the same chemical-taxon pair are likely to appear in both the training and test set. We call this data splitting *totally random*.

#### 2.5.2 Splits by chemical compound

Stratification by chemical compound ensures that chemicals are not shared between training and test set. For the split *by occurrence of chemical compounds*, compounds are sorted by the number of experiments performed on them, *i*.*e*., those with most experiments at the top. Then, the first five compounds are put into the training set and the sixth is put into the test set. This is repeated with the subsequent compounds until all are distributed. The five cross-validation folds are filled accordingly, *i*.*e*., the most common compound goes to fold 1, the second most common to fold 2, and so on. However, with this split, it is still likely that similar chemicals are shared between the training and test set, and between the cross-validation folds.

In ADORE, training and test splits as well as cross-validations folds for both splitting schemes are available. Since the split by occurrence of chemical compounds puts one sixth, *i*.*e*., 17%, of the data points in the test set, the associated ratio of 83:17 was maintained for the totally random split to have comparable sizes across the data subsets.

## 3 Methods

In this section, we introduce the regression models applied to the dataset using the log10-transformed mass and molar LC50 as response values.

All code was written in Python 3 using established ML libraries. It is available in a public repository^*^ where the package versions are specified in the environment.yml file.

### 3.1 Models

We compared four established regression techniques that can be applied to QSAR. Least Absolute Shrinkage and Selection Operator (LASSO) is a linear regression technique with inherent feature selection. The tree-based models random forest and eXtreme Gradient Boosting (XGBoost) are commonly used in ecotoxicology and can be considered state-of-the-art. Gaussian process regression is more complex and computation-intensive but has the advantage to provide uncertainty estimates. Random forest and LASSO models were developed using scikit-learn ^32^, XG-Boost models with the XGBoost package ^33^, and the GP models were built using the gpflow package ^34^. The LASSO has the lowest computational cost of the considered models. The RF models take shorter to run than XGBoost models. The hyperparameters of each model are summarized in the supplemental Table 2.

#### 3.1.1 LASSO

The LASSO is a regularized linear regression model. Regularization introduces a term to the loss function of ordinary least squares (OLS) regression that favors smaller regression coefficients. For LASSO, regularization shrinks coefficients to zero and is therefore performing inherent feature selection as only features with non-zero coefficients are retained. In the closely related Ridge regression, coefficients are shrunk towards zero but do not reach zero. The importance of the regularization term is determined using the regularization coefficient, *α* (alpha), which is the only hyperparameter of LASSO.

We employed a two-step procedure that has the advantage to give a smaller set of selected features than directly using the results from LASSO ^35^. In the first step, the LASSO was fit on the training data for a range of the hyperparameter *α*. For each *α*, all features with non-zero coefficients were retained. In the second step, a Ridge regression model is trained using only the non-zero coefficients (if there are any), and then evaluated on the validation data. The *α* with the best validation error is selected.

#### 3.1.2 Random forest

Random forest is an ensemble learning method using decision trees that constructs mutually-independent trees using the response value as target variable. Each tree is learned on a bootstrap sample of the training data, a procedure known as bootstrap aggregation (“bagging”) ^36^. Trees are further de-correlated by only considering a subset of features for each split. For regression, the results of each tree are averaged to obtain the prediction. Typically, a few hundred trees are learned with depths in the range of a few dozen to a few hundreds. We optimized the following hyperparameters: number of trees (n_estimators), maximum depth of a tree (max_depth), minimum number of samples required to split an internal node (min_samples_split), number of bootstrap samples (max_samples), and number of features when looking for the best split (max_features).

#### 3.1.3 XGBoost

Gradient boosting is another ensemble learning technique that, in contrast to the bagging approach of RFs, develops models sequentially using the error of the predecessor model as target variable. In the case of regression, the residuals, *i*.*e*., the difference between the true and the predicted value, are minimized. Gradient boosting has been refined in the extreme gradient boosting algorithm ^33^, which is more scalable than gradient boosting and the state-of-the-art implementation of boosted decision trees. Typically, XGBoost trees are less deep than RF trees, the depth ranging up to a dozen nodes. We optimized the following hyperparameters: number of trees (n_estimators), shrinkage of step size (eta), minimum reduction of loss to split a node (gamma), maximum depth of a tree (max_depth), minimum weight of a child node (min_child_weight), and subsample ratio (subsample).

#### 3.1.4 Gaussian process regression

Gaussian processes are state-of-the-art Bayesian tools for regression ^37^, classification ^38^, and dimensionality reduction ^39^. A GP for linear regression uses a Gaussian prior over the weights of the regressor. It couples them with a least square error loss function (Gaussian likelihood), which allows for computing in closed form the best prediction for each input and its confidence interval. By relying on the kernel trick ^37^, GP can also solve nonlinear regression problems in closed form. It is the main feature of GP to provide accurate predictions, which naturally come with confidence intervals. On the other side, GP come with high computational complexity (*i*.*e*., cubic in the number of samples), which renders them the slowest model we compare. See appendix A.1 for details on the GP implementation.

### 3.2 Hyperparameter optimization

For each combination of model, molecular representation, data splitting scheme, and concentration type, we chose the corresponding optimal hyperparameter(s) using gridsearch. For each hyperparameter setting, 5-fold cross-validation on the training data was employed and the hyperparameter setting with the lowest cross-validated root mean square error (RMSE) was selected. Then, the model with the best cross-validation performance based on RMSE was retrained on the entire training set and evaluated on the test set. We report both cross-validation and test error.

### 3.3 Metrics

To evaluate the cross-validation and test runs, we calculated micro-average RMSE, mean absolute error (MAE), and the coefficient of determination *R*^2^ (see Appendix B.1). For the test runs, we also evaluated macro-averaged metrics (see Appendix B.2).

In contrast to *R*^2^, RMSE and MAE have the same dimension as the response, the log10-transformed LC50. Accordingly, an RMSE or MAE of 1 translates to one step on the log10 scale, *i*.*e*., one order of magnitude on the original, non-transformed, scale. This direct relation to the response unit allows for an intuitive interpretation of error values.

### 3.4 Feature importance

For the tree-based models RF and XGBoost, we investigated two types of feature importances: permutation based feature importances, calculated using the scikit-learn function sklearn.inspection.permutation_importance, and SHAP (SHapley Additive exPlanations) values ^40^. Feature importance methods can be distinguished by their scope, *e*.*g*., do they provide feedback on the entire model (global scope), or do they explain an individual prediction (local scope) ^41^. The permutation feature importance measures the increase in prediction error when permuting the values of features, providing global information about the model. On the other hand, SHAP values are a local method as they are calculated for individual predictions. They can be averaged for a global interpretation of the model.

### 3.5 Reporting

Several best practices for ML-based science and QSARs have been proposed. We evaluated our work against three checklist-based reporting schemes: (1) the REFORMS checklist for ML-based science ^22^, (2) potential pitfalls related to data leakage ^15^, and (3) a QSAR-specific checklist ^18^, which has been extended to the application of ML to QSARs ^19^. We consider our approach to go beyond QSAR through the integration of species-specific and experimental data. Nonetheless, these guidances are still relevant to our work. The detailed self-assessments can be found in the supplementary information.

## 4 Results and Discussion

### 4.1 Data quality & variability

Data is the basis for every model. *Ipso facto*, model performance is limited by the quality of the data it was trained on. Reliable predictions can only be obtained within the range of data (*i*.*e*., range of toxicity and range of features) according to the boundingbox approach, a simple applicability domain technique. Fig. 2 shows the training and test set distribution of the response value (LC50 in *mol/L*) and three relevant chemical features: molecular weight, water solubility, and logP. Training and test set were constructed to cover a similar range of the response values as well as the chemical properties.

**Fig. 2.**
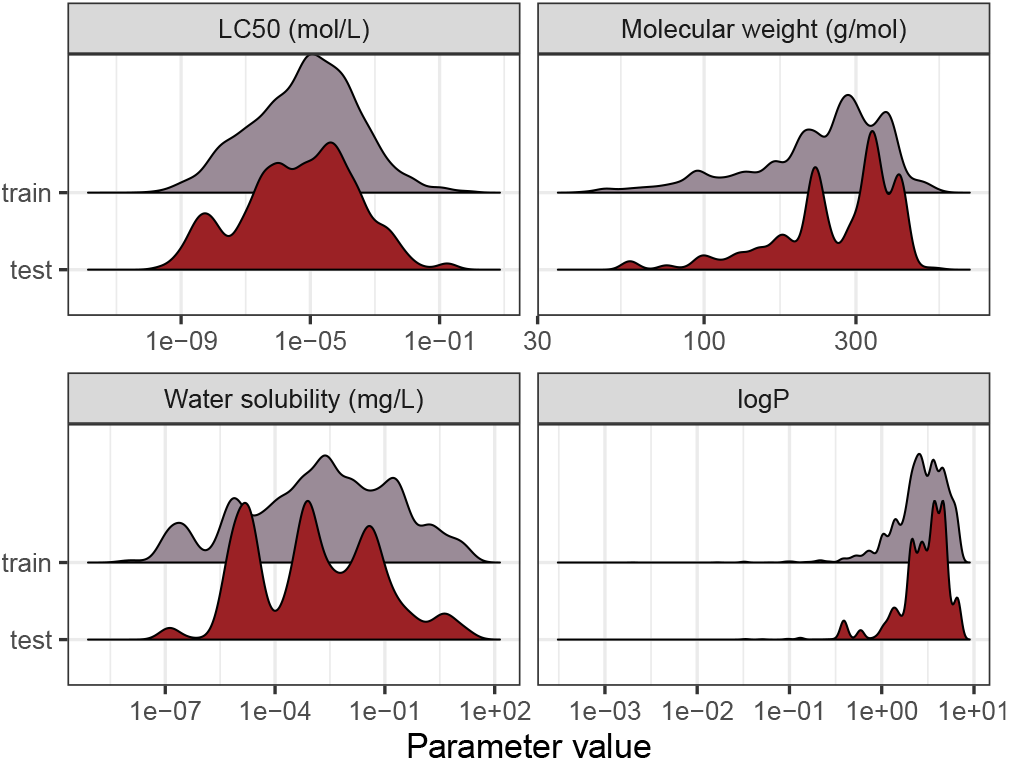
Distribution of the training and test set from the split by occurrence of chemical compounds for LC50 (in *mol/L*), molecular weight (in *g/mol*), water solubility (in *mg/L*), and logP.

ADORE contains repeated experiments that do not necessarily share the LC50 value. Most experiments have only one or a few values associated with them (Fig. 3(A)). Nonetheless, the LC50 values can vary over several orders of magnitude, as is depicted in Fig. 3(B) for fish tests repeated at least 25 times. *In vivo* data, by default, is highly variable, even within strictly standardized experimental settings such as the OECD TG 203^2^.

**Fig. 3.**
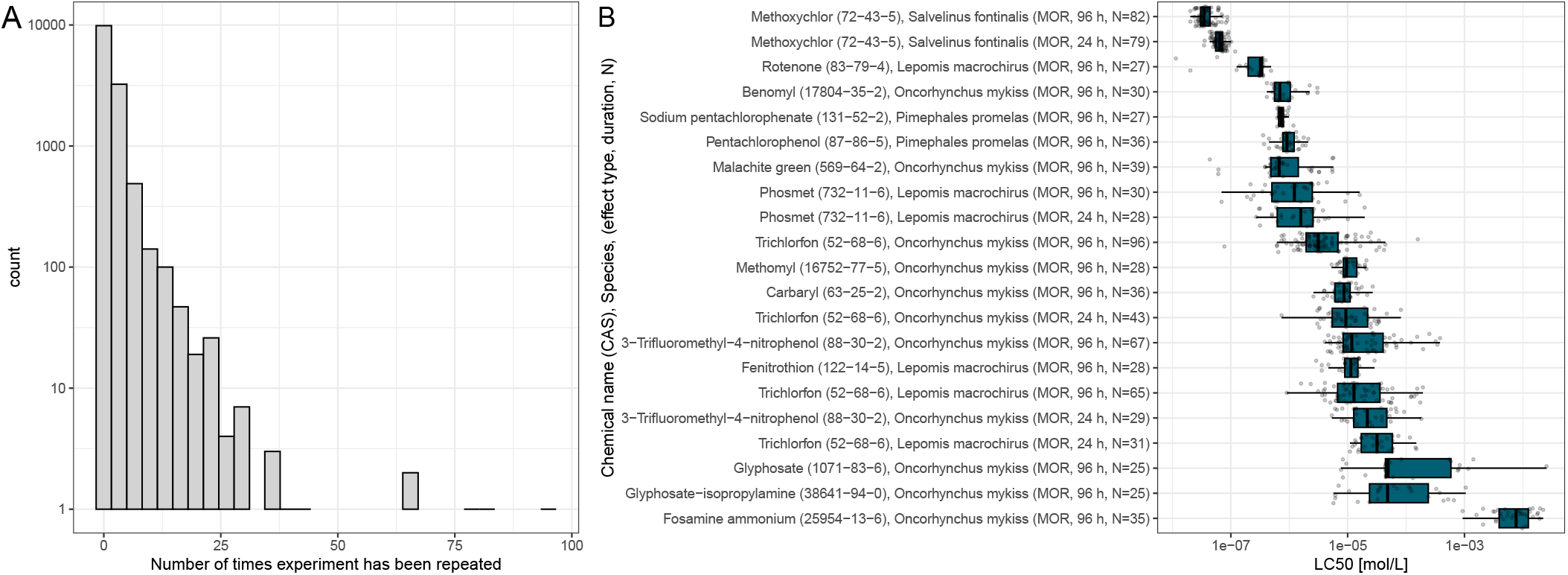
A: Histogram of the number of data points associated with a combination of chemical, species, and experimental conditions (*i*.*e*., media type, observation duration, concentration type, and exposure type). B: Boxplot of toxicity values for experimental conditions with at least 25 values. Y-axis labels indicate the chemical name, CAS number, the species it was tested on, the effect group, observation duration, and the number of data points. For fish, all tests were carried out for the effect group mortality (MOR).

### 4.2 Modeling results

#### 4.2.1 Validation results

Here, we discuss cross-validation results. The results on the test set are described in Sec. 4.2.2.

##### Data splitting scheme

For the totally random split, we achieve much better performances than for the split by occurrence, independent of concentration type, model, and molecular representation (see Fig. 4 and supplemental Figures 1 and 2). For models trained using a molecular representation, the RMSE does not exceed 0.90, MAE does not exceed 0.65, and *R*^2^ is above 0.65, for all combinations. For the tree-based models, RF and XGBoost, the RMSE is around 0.50, MAE around 0.30, and *R*^2^ is reaching 0.90. Despite having been achieved on the same dataset, these performances are substantially better compared to the split by occurrence. This shows how data leakage produces artificially inflated performances.

**Fig. 4.**
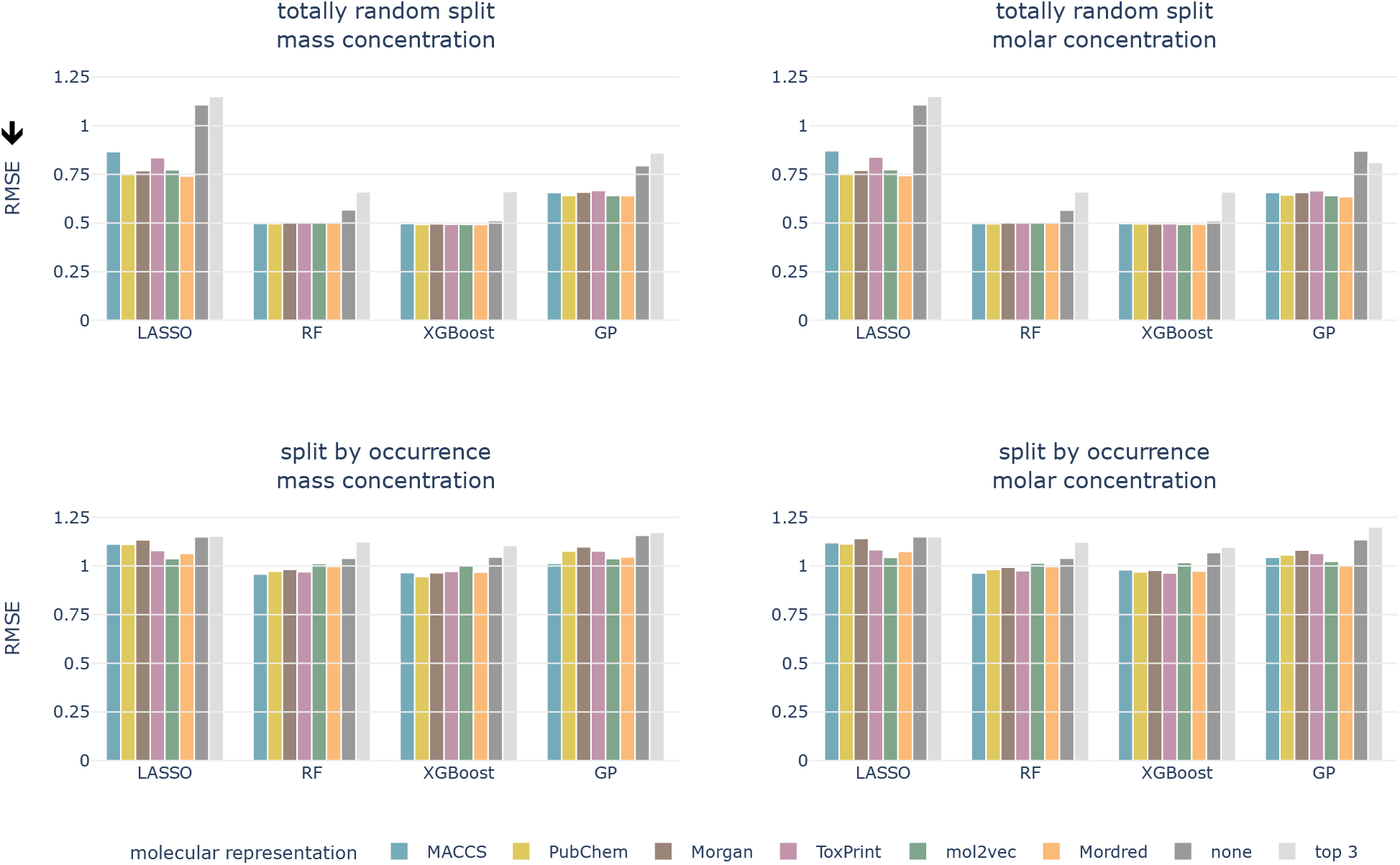
Cross-validated RMSE for both data splittings, concentration types, all models, and molecular representations. Arrow indicates the lower the better.

Signatures of data leakage can also be seen in the selected hyperparameters. For LASSO, the regularization parameter *α* is consistently smaller for the totally random split (0.00001 – 0.00025) than for the occurrence split (0.00631 – 0.10000) (supplemental Table 3). A smaller *α*, relates to more features being added to the model, which can be interpreted as the model attempting to memorize the training data. We observe the same behavior for the tree-based models, but less consistently. For the RF models, more and deeper trees are selected (supplemental Table 4) for the totally random split. Also for the XGBoost models, deeper trees are grown for the totally random split than for the occurrence split (supplemental Table 5). Deep trees can be related to overfitting.

##### Concentration type

The models perform on par for both the log10-transformed mass and molar LC50 independent of the data splitting scheme and the molecular representation.

##### Model

The tree-based models perform best for all combinations of data splitting schemes and concentration type, followed by GP regression. The linear model, LASSO, performs worst.

##### Molecular representation

The six representations perform similarly for all combinations of concentration type, data split, and model, shown as colored bars in Fig. 4 and supplemental Figures 1 and 2. Additional bars indicate performances with only experimental, chemical and taxonomic properties (‘none’, *i*.*e*. no molecular representation) or using only three chemical properties (‘top 3’).

For the remainder of the study, we focus on the split by occurrence of chemical compounds, since it reduces the risk of data leakage compared to the totally random split, and on the molar-based LC50, since it more closely resembles the outcome of toxicity tests. For the occurrence split, all combinations of concentration types, models, and molecular representations achieve an RMSE of around 1, which means that, globally, the LC50 can be predicted within an order of magnitude (see bottom row of Fig. 4). For the moment, we do not restrict ourselves to a molecular representation but first evaluate the test performance.

#### 4.2.2 Performance on test set

We evaluated the best models on the test set for molar LC50 and the split by occurrence of chemical compounds. Test and crossvalidation (micro-average) RMSE and *R*^2^ are shown in Fig. 5. The test performance is comparable to the cross-validation performance, *e*.*g*., the tree-based models perform better than GP and LASSO, and for models trained on a molecular representation, the RMSE varies around 1.0 and *R*^2^ around 0.6.

**Fig. 5.**
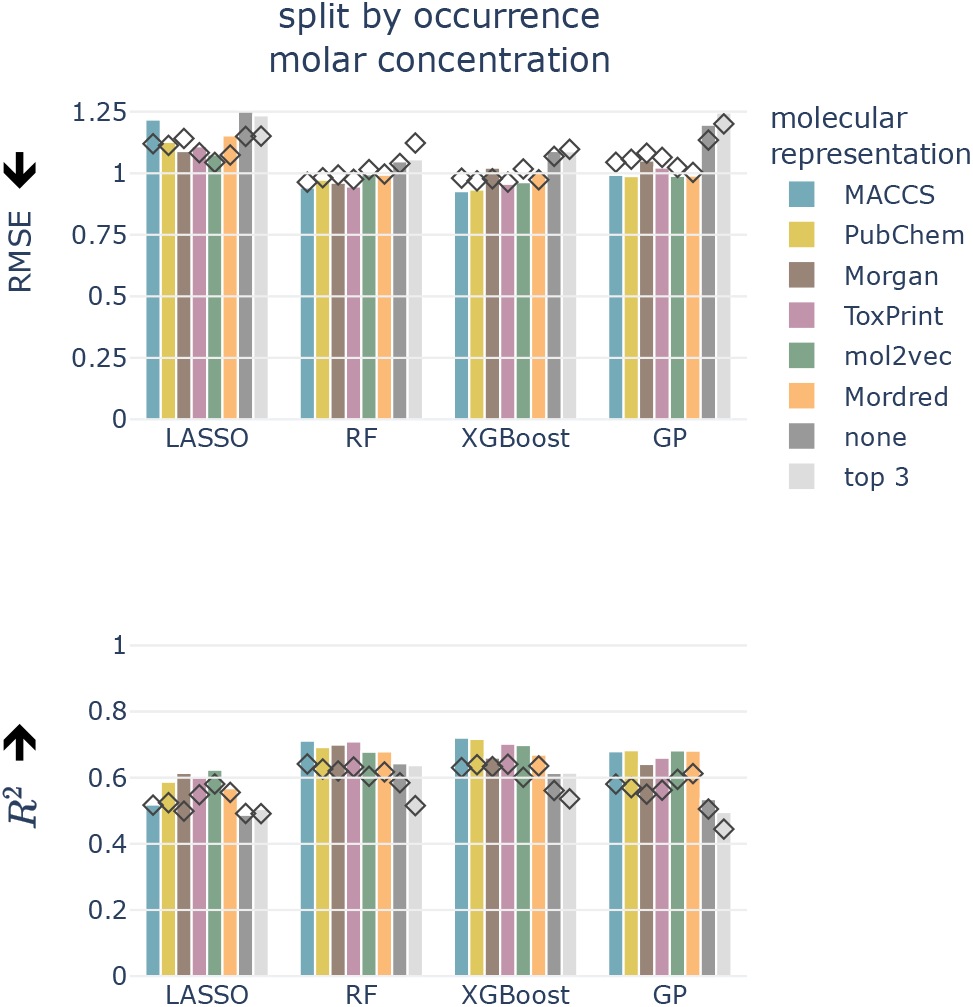
Test (depicted as bars) and cross-validated (diamonds) RMSE and *R*^2^ for molar LC50, split by occurrence of chemical compounds, all models, and molecular representations. Arrows indicate the lower/higher the better.

Also for the test set, the six molecular representations perform on par for each model. This indicates that these molecular representations, in combination with the chemical properties (with the exception of Mordred, since this is a combination of molecular representation and chemical properties), are equally valid descriptors of the underlying chemical space. The molecular representations are necessary features as models without them perform worse.

The macro-averaged RMSEs only show minor differences compared to the micro-averaged RMSE. Tree-based models perform best independent of the average type. According to supplemental Fig. 3, the best micro-averaged test performance is achieved with an XGBoost model trained on MACCS fingerprint features (*RMSE*_*µ*_ = 0.927). The macro-averages for chemicals and taxa combined and for taxa only are also best for the MACCS fingerprint (RF, *RMSE*_M_ = 0.904 and XGBoost, *RMSE*_T_ = 0.938, respectively). The chemical macro-average is best for RF and ToxPrint (*RMSE*_C_ = 0.845) (supplemental Fig. 3).

### 4.3 Including phylogenetic distances

For GP, the phylogenetic pairwise distances can be used for modeling by adding a pairwise distance kernel. This does not improve the predictions, as GP models with and without pairwise distances perform similarly, both during cross-validation (supplemental Fig. 4) and when testing (supplemental Fig. 5).

### 4.4 Explainability

Machine learning models are widely considered as black box models, where the prediction process is mostly opaque. However, there exist several approaches that render models more explainable and allow to better understand the relevance of input features.

#### 4.4.1 Residuals

Residuals can aid in identifying correlations between features and local model performance. A residual is the difference between the predicted and the true value. A negative residual corresponds to an overprediction of the toxicity, *i*.*e*., the chemical was predicted more toxic than it actually is, while a positive residual is an underprediction of the toxicity. Given the goal of chemical hazard assessment, the latter is the more problematic case.

To get an intuition about the variation of predictive power across the toxicity range, we analyzed the correlation between residuals and true LC50 values (Fig. 6(a)). Lower LC50 values (corresponding to higher toxicity) are correlated with higher residuals indicating that these values get underpredicted. Also, higher LC50 values (corresponding to lower toxicity) are correlated with lower residuals indicating that these get overpredicted. This phenomenon is also known as “regression to the mean”.

**Fig. 6.**
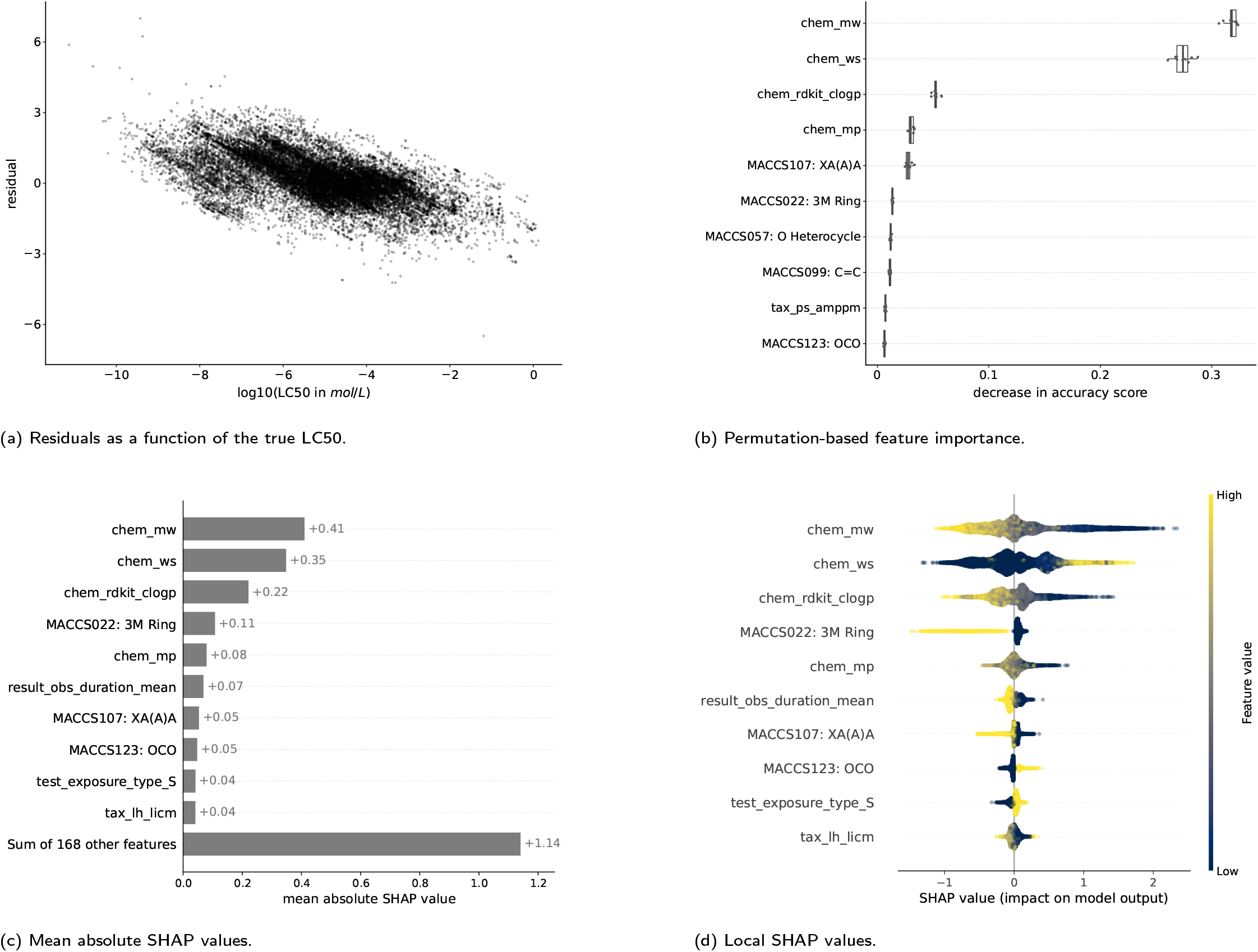
Feature importances for the XGBoost model trained with the MACCS fingerprint to predict molar LC50. For both methods, the respective top 10 features are shown. The features are listed in the supplemental Table 1. For chemical properties, the prefix is “chem”, for experimental features, “result” or “test”, for taxonomic properties, “tax”. The bits from the MACCS fingerprint contain the bit number and the corresponding SMARTS string.

The stripes in the plot correspond to repeated experiments with varying experimental outcomes but the same prediction. The variability of repeated experiments, visualized in Fig. 3, cannot be captured by the models as repeated experiments have exactly the same feature values for chemical, taxonomic, and experimental properties.

#### 4.4.2 Feature importance

Given the inconsistent predictive capacity of models on a local level (*i*.*e*., the difference between the ground truth and the predicted value), we abstain from definitive conclusions regarding feature importance. Nevertheless, including feature importance contributes to the discussions between regulators, ecotoxicologists, and data scientists, on the explainability of models and the role of the currently available feature importance methods.

Here, we show the feature importances for the XGBoost model trained with the MACCS fingerprint to predict molar LC50. The results for other combinations of models and molecular representations are similar and for most combinations, three chemical properties contribute most according to both feature importance methods. Molecular weight (*chem_mw*), water solubility (*chem_ws*), and logP (*chem_rdkit_clogp*) have the highest importance by a large margin compared to the next features (Fig. 6(b),(c)). Nevertheless, the other 168 features still explain more (+1.14) than the three top features together (0.41 + 0.35 + 0.22 = 0.98) (Fig. 6(c)). This is confirmed by the model runs on the top 3 features only (light gray bars in Figs. 4 and 5), which perform worse than models with chemical, taxonomic, and experimental features. The SHAP values by data point (Fig. 6(d)) allow for an interpretation of how higher or lower values of a property are correlated with higher or lower toxicity predictions. Positive SHAP values correspond to a higher LC50 and, thus, lower toxicity. As an example, high logP values, which are corresponding to increased lipophilicity, lead to negative SHAP values. This trend is inversed for water solubility. These observations are consistent with the ecotoxicological principle that compounds with higher lipophilicity will accumulate more in fatty tissues and, as an effect, elicit higher toxicity. As is intuitive, a longer observation duration (*result_obs_duration_mean*) leads to lower SHAP values and therefore higher toxicity. A higher molecular weight leads to higher toxicity, which is likely correlated to larger molecules also being more lipophilic ^42^ (supplemental Fig. 12). For other features, there is no straightforward interpretation: The binary coded static exposure (*test_exposure_type_S*), *i*.*e*., 1 for static exposure and 0 for other exposure types, shows higher SHAP values for the static exposure (high value on the binary scale).

As an additional aspect, there are only two taxonomic features among the most important features, the DEB parameter *tax_ps_amppm* for the permutation-based feature importance and the ultimate body length (*tax_lh_licm*) for SHAP. This indicates that the provided species-related features do not enable the models to adequately capture the sensitivity profiles of the chemicals.

#### 4.4.3 Species sensitivity

Species sensitivity distributions (SSDs) are a common method in ecotoxicology that integrate toxicity data of several species for one chemical (the suggested minimum number of species is 15) ^43^. It serves to identify the percentage of tested species at risk at a certain concentration. Decades after the introduction, SSDs are still the subject of active research ^44,45^. Here, we produced SSDs for compounds that have been tested on at least 15 different fish species to investigate how well the model predictions match the species sensitivity of the original biological data. The sensitivity of species to a chemical can span several orders of magnitude while the range covered by the model predictions is far smaller and does not follow the sigmoidal shape of the ground truth as can be seen for four pesticides, a contraceptive compound, and potassium cyanide in Fig. 7. We therefore conclude that species-specific sensitivities are not adequately distinguished by our models.

**Fig. 7.**
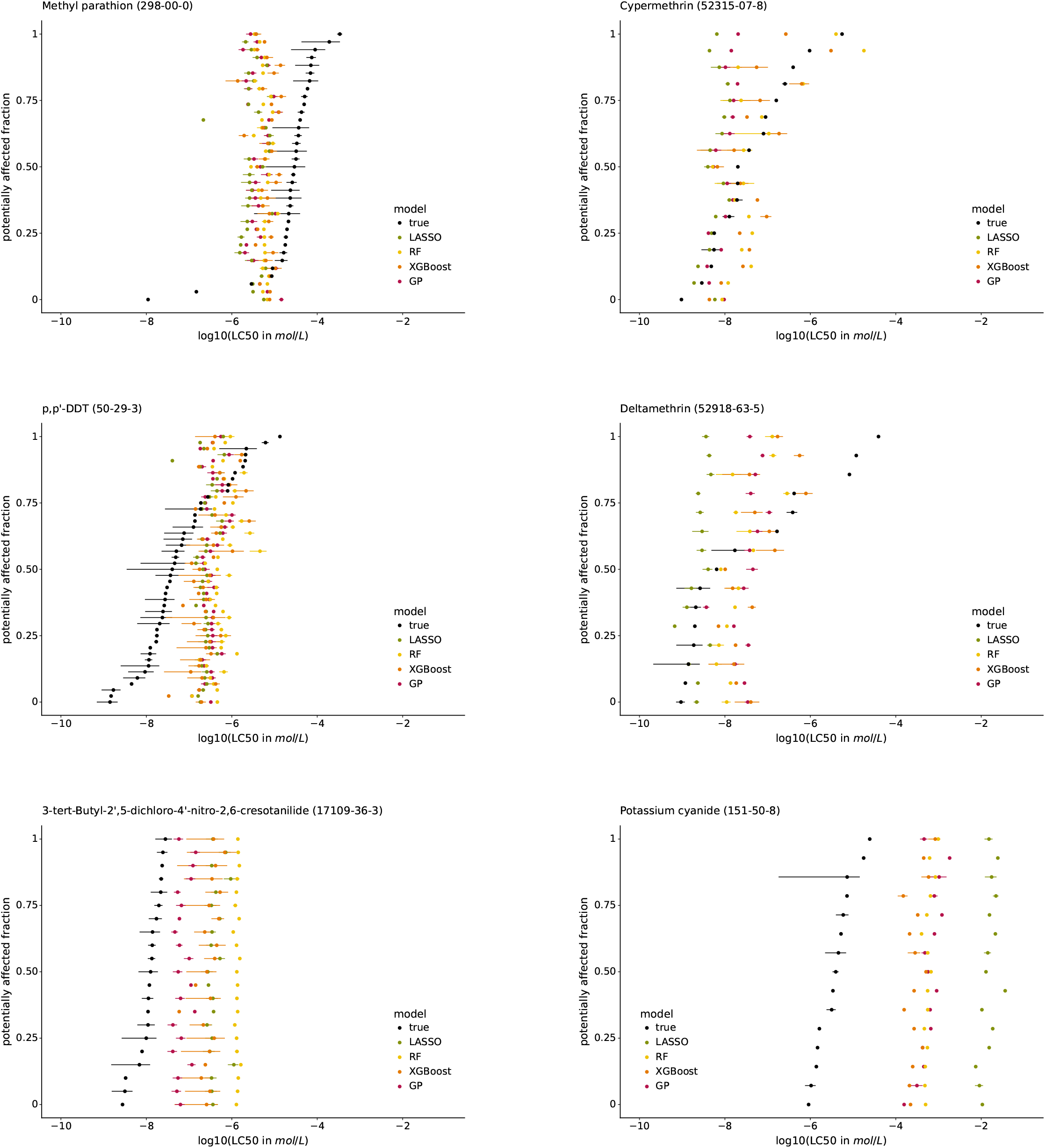
Species sensitivity distributions (SSDs) of the pesticides methyl-parathion, cypermethrin, p,p’-DDT, and deltamethrin, the contraceptive compound 3-tert-Butyl-2’,5-dichloro-4’-nitro-2,6-cresotanilide, and potassium cyanide. The species are sorted according to their median true LC50. The SSDs of the other chemicals tested on at least 15 species are in the supplementary material.

### 4.5 Reporting: Self-evaluation

We are in line with the REFORMS reporting standards ^22^ by using a published benchmark dataset and by making all code and data openly available.

We are confident to have considered all points from the data leakage and reproducibility checklist by Kapoor *et al*. ^15^. Apparent shortcomings stem from the dataset itself and are not straightforward to evaluate, *e*.*g*., a potential sampling bias in the test distribution. We compared the distributions of key features between training and test sets, but cannot definitively exclude the possibility of a non-obvious bias. We are transparent about other potential pitfalls inherent to the dataset, here and in Ref. ^24^, where we present the dataset and its curation.

According to the guidance documents by Cronin *et al*. and Belfield *et al*., the highest uncertainty in our work is related to the quality of the original biological data, since we did not verify all data points against the original literature. Likewise, measured chemical concentrations both in the exposure medium and internal concentrations of the organisms are not included in the data. Additionally, acute mortality is an unspecific endpoint, as it only accounts for the death of a specimen. Chemicals can cause death through a number of different mechanisms, which are not distinguished in the effects records. This leads to high uncertainty on the mechanistic interpretability of this data.

Overall, our work reflects awareness of all raised concerns. We openly communicate the drawback of the skewed dataset, which does not allow to split the data according to chemical and species at the same time. We consider it more important to avoid data leakage related to chemicals than to the species, since the latter would be counter-intuitive to the ecotoxicological principle of using model/surrogate species.

### 4.6 Relevance for environmental hazard assessment

Determining toxicity is a routine part of the regulatory framework to ensure the safety of chemicals on the market. The integration of *in silico* methods into this framework is widely discussed ^4,46–49^. Reliable computational tools could predict toxicological outcomes of chemicals to reduce the need for animal testing. More importantly, they could serve as pre-selection tools to find candidate molecules for a use case in accordance with safety requirements. This would move chemical design and production closer to the safe and sustainable by design (SSbD) principle, ensuring chemical safety already during the design phase ^50^. To be fit for this purpose, model predictions should be consistent and explainable across a broad chemical and taxonomic space. Here, regulators need to lead the way by, for example, specifying the expectations on NAMs in general and *in silico* methods in particular, such that they can be included into an updated paradigm for regulatory hazard assessment ^51^.

The global performance of our models is satisfactory, as the LC50 could be predicted within one order of magnitude. However, on a local level, model performance mainly depends on the chemical properties. Also, the species-specific features are not sufficiently informative to explain species differences. The toxicity of many chemicals is either over– or under-predicted, but not in a consistent manner. If the chemicals were generally predicted to be more toxic than they are, this consistently conservative estimate would be in line with the precautionary principle and could be implemented into a regulatory workflow as a pre-selection step. However, the development of acceptable NAMs for the regulatory framework is an iterative process requiring concerted stakeholder efforts when refining model requirements and performances. Accordingly, scientists and regulators need to be closely connected to ensure persistent progress towards this shared goal. We strongly believe in the potential of ML methods to be an asset in this process. In the following subsection, we point out several routes that could, from the modeling perspective, lead to further improvements in both performance and consistency.

### 4.7 Limitations & future work

Some limitations are related to the underlying data, while others are of technical or conceptual nature.

The ADORE “t-F2F” dataset contains results from tests performed on over 100 species for almost 2,000 chemicals. Since the regulatory use case is focused on few surrogate species, the use of such a broad dataset has drawbacks. The OECD TG 203^2^ for fish acute mortality suggests six model species as surrogates, which renders models trained on single species data, such as the ADORE challenges “s-F2F-1”, “s-F2F-2”, and “s-F2F-3”, closer to the regulatory use case than models based on the “t-F2F” challenge. The chemicals might not be represented adequately, *e*.*g*., there might be a better descriptor than the currently used molecular representations. Also, the chemicals are described based on canonical SMILES, which do not capture isomerism and 3D structures. Additionally, the applicability domain is only partly defined. Other approaches might help to better understand the underlying chemical space, providing information on which additional data could prove useful in future work. Regarding the experimental data, there is only information on the use of active ingredients in isolation or formulations (*test_conc1_type*), and not on the composition of these formulations and the percentage of active ingredient contained in them.

Biologically, by choosing acute mortality, we opted for an unspecific effect not linked to specific modes of action. By refining the scope of effects to either groups of chemicals with specific mechanisms or to effects that are closely coupled to specific modes of action, better model performances could be expected. On the other side, given far less training data, this could also lead to worse model performances or overfitting. Exploring the application to other levels of biological organization is a worthy goal, despite acute mortality being one of the most significant ecotoxicological effects within the current regulatory framework ^52^. Feature importance analysis in conjunction with species sensitivity distributions indicated that the current taxonomic features are not sufficiently capturing species-specific differences. Zubrod *et al*. (2023) ^7^ expanded their feature space with more species-specific features at the cost of a smaller species coverage, which is a focus on few, well-covered (surrogate) species representing a trade-off worth to explore. Likewise, efforts exist to map the conservation of potential molecular targets for toxic effects across species using genetic data. However, given the low specificity of acute mortality, this is currently unlikely to be adapted ^53^. Future work could include phylogenetic distances as a feature on a superficial level, *e*.*g*., by using the phylogenetic distance to a single reference species instead of using the complete pairwise distance matrix.

Apart from these data and technology related limitations, other model types, such as the pairwise recommender systems deployed by Viljanen *et al*. (2023) ^54^, could be explored.

On a broader level, this study is based on *in vivo* data and aimed to assess the suitability of ML as an alternative to animal experiments. Meanwhile, other alternative methods have been established or are under development, for example, tests based on early life stages (fish embryo acute toxicity test; OECD TG 236^55^) and on isolated cell cultures (*in vitro*, fish cell line acute toxicity assay; OECD TG 249^56^). These NAMs have a high potential as reliable and reproducible data sources, which can be used to train models for potentially higher predictive performance, reducing the reliance on *in vivo* data. They may also help in filling data gaps on a local level. The integration of multiple alternative endpoints through ML into a toolbox-based framework may benefit the regulatory process, compared to evaluating individual NAMs against the currently accepted endpoint fish acute mortality. As described earlier, we believe that this effort needs to be undertaken in close collaboration between researchers and regulators to cater to the strengths of the individual methods while ensuring both public and environmental safety.

### 4.8 Comparison with previous studies

Several studies have applied ML regression models to predict ecotoxicological outcomes in fish. Since results were obtained from data with different taxonomic (single-species vs across-species and across-taxa vs multiple species) and chemical scopes, comparison is difficult, both among the previous studies and to our work. Comparison is additionally hindered by different train-test-splitting schemes. This substantiates the necessity of adopting the use of benchmark datasets and best practices for ML-based research and its dissemination going forward.

Similar to us, Zubrod *et al*. (2023) modeled multiple species simultaneously and included species-specific data from the Add my Pet database in addition to chemical properties and the molecular representations ToxPrint and Mordred to predict log10-transformed mass LC50^7^. However, their chemical space was limited to pesticides. Their freshwater fish dataset, containing 1,892 samples from 92 species and 360 pesticides, was obtained from the Envirotox database ^57^, which largely overlaps with ECO-TOX. For a species-chemical combination, they averaged all data points. They performed random data splitting, which corresponds to our totally random split, and splitting stratified by toxicity values. No stratification by chemical and/or species was mentioned. They trained RF models using 10-fold cross-validation with varying feature sets and obtained test RMSE values of 0.54, which is comparable to our results from the totally random split. According to mean absolute SHAP values, water solubility and logP are the most important predictors in their final model.

Additional works, often focusing on the “aquatic triad” of algae, crustaceans, and fish, which are commonly used as surrogates for the different trophic levels in ecotoxicology, are discussed in Appendix C.

## 5 Conclusions

Our study focused on the implementation and interpretation of machine learning methods to predict fish acute mortality. We trained four types of models to predict the lethal concentration 50 (LC50) of 1,905 compounds on 140 fish species. We found that tree-based models, specifically RF and XGBoost, performed best and were able to predict the log10-transformed LC50 with a root mean square error of 0.90, which corresponds to (slightly less than) an order of magnitude on the original LC50 scale. However, on a local level, the models are not yet accurate enough to consistently predict the toxicity of single chemicals across the taxonomic space. The models were found to be mainly influenced by a few chemical properties and to not capture taxonomic traits, and thus species-specific sensitivities, sufficiently.

In conclusion, while ML models show promise in predicting fish acute mortality, there are still limitations that need to be addressed. We see this study as a contribution to the ongoing discussions on how machine learning can be integrated into the regulatory process, while further research and improvements are needed to achieve better explainability and, as a result, foster acceptance. To progress the field as a whole beyond individual studies, transparency, comparability, and reproducibility need to be considered in the development of models.

## Supporting information

Supplementary Material – Self-assessment

## Code and Data Availability

The code is available on https://renkulab.io/gitlab/mltox/mltox-model. The ADORE dataset is available on ERIC, the institutional data repository of Eawag (https://doi.org/10.25678/0008C9) and in the repository https://renkulab.io/gitlab/mltox/adore. The modeling repository https://renkulab.io/gitlab/mltox/adore-modeling contains code on how to load the data, prepare it for modeling, *e*.*g*., create one-hot and multi-hot-encodings for categorical features, apply the train-test-split for 5-fold cross-validation, and train and evaluate RF models.

### Author Contributions

LG and CS share first authorship of this work.

LG: Visualization; Software; Investigation; Formal Analysis; Data curation; Conceptualization; Writing – original draft; Methodology; Project administration; Validation; CS: Visualization; Software; Investigation; Formal Analysis; Data curation; Conceptualization; Writing – original draft; Methodology; Project administration; FPC: Conceptualization; Writing – review & editing; Resources; Methodology; Project administration; Supervision; KS: Conceptualization; Writing – review & editing; Funding acquisition; Methodology; Project administration; Supervision; MBJ: Conceptualization; Writing – original draft; Writing – review & editing; Funding acquisition; Methodology; Project administration; Supervision

## Acknowledgements

We thank Guillaume Obozinski from the Swiss Data Science Center for valuable discussions and input. This work was made possible through the SDSC grant “Enhancing Toxicological Testing through Machine Learning” (project No C20-04) and partly carried out in the framework of the European Partnership for the Assessment of Risks from Chemicals (PARC) and has received funding from the European Union’s Horizon Europe research and innovation program under Grant Agreement No 101057014. Views and opinions expressed are however those of the authors only and do not necessarily reflect those of the European Union or the Health and Digital Executive Agency. Neither the European Union nor the granting authority can be held responsible for them. The graphical abstract was created with BioRender.com.

## A Models

### A.1 Details on the Gaussian process implementation

The Gaussian process learns from the similarity of the data points that are presented to it through a kernel function. The kernel function is calculated for each pair of data points leading to an *n × n* symmetric matrix, where *n* is the number of data points, and each entry corresponds to the similarity of two data points. We propose an additive kernel that separates the different groups of variables:

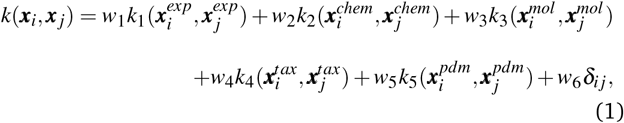

where *w*_*i*_ is the relative strength of each kernel and *k*_*i*_(***x***_*i*_, ***x*** _*j*_) is the well-known squared exponential (SE) kernel

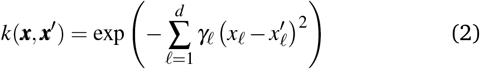

for the experimental features (exp), chemical properties (chem), molecular representation (mol), and taxonomic properties (tax). For the taxonomic pairwise distances (pdm), we used a pairwise distance kernel, which has the pairwise phylogenetic distance of the two species associated with the respective data points as each entry.

The SE kernel can be used in an unweighted fashion with the same lengthscale *γ*_1_ = *γ*_2_ = … = *γ*_*d*_ = *γ* for all *d* features. Alternatively, a characteristic lengthscale *γ*_*ℓ*_ per feature is optimized using a procedure called automatic relevance determination (ARD).\ The inverse of the characteristic lengthscale determines the relevance of each feature ^37^. The SE kernels for the first four groups of features were used with ARD and initialized with *w*_*i*_ = 1 and *γ*_*iℓ*_ = 3.

To substantially reduce computation time, we used a sparse GP regression algorithm ^58^, which is implemented in the gpflow package ^34^ in the function gpflow.models.SGPR. The compute time for the 22k training entries could be reduced from more than a day to a few hours. The sparse approach constructs an approximation using a small set, typically a few hundred, of inducing points, which are representative points capturing the data structure. The number of inducing points is the only hyperparameter for the GP model. We selected the inducing points using k-means clustering, where k corresponds to the number of inducing points. The clustering algorithm, implemented using scikit-learn, returns the cluster centers, and not actual data points, as input for the sparse GP regression.

See Rasmussen and Williams ^37^ and the appendix of Gasser *et al*. ^59^ for a more detailed description of Gaussian processes.

## B Metrics

< this section has been moved from the methods section >

### B.1 Micro-averaged metrics

For cross-validation and testing, we calculated the micro-average RMSE, MAE, and *R*^2^,\

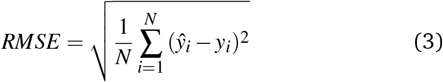

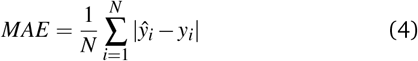

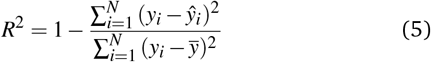

of the respective data subsets containing *N* samples. We call *y*_*i*_ the measured response and *ŷ*_*i*_ the predicted response for entry *i. y* is the average response in the respective data subset, *e*.*g*., the test data if we calculate test *R*^2^. In micro-averaged metrics, each data point has the same weight. This means, for example, that chemicals appearing more often will be over-represented.

We do not report the squared Pearson coefficient, *r*^2^, that is used in older QSAR studies, as it is not an appropriate metric in our case. When dealing with nonlinear models, *r*^2^ ≠ *R*^2^, and model selection based on the *r*^2^ can lead to a wrong interpretation of the results ^60,61^. In fact, not only the squared Pearson coefficient treats positive and negative correlations equally, but also a perfect correlation (*r* = 1) does not necessarily imply that *y*_*i*_ = *ŷ*_*i*_ for every *i* (see Khandelwal ^62^ for a didactic explanation).

### B.2 Macro-averaged test metrics

For the test sets, we also calculated macro-averaged test metrics to account for repeated experiments on chemicals and species. These give the same weight to chemicals and/or taxa instead of micro-averaged metrics that give the same weight to individual data points. A test set containing *N* samples with repeated experiments has *N*_*C*_ *< N* chemicals and *N*_*T*_ *< N* taxa. We use *c* to indicate a chemical and *t* to indicate a taxon. A repeated experiment (*i*.*e*., a (*c, t*) couple) has *n*_*ct*_ instances. We call *n*_*c*_ the number of times chemical *c* was tested and *n*_*t*_ the number of times that taxon *t* was tested.

#### Micro-averaged root mean square error (*µ*RMSE)

This metric corresponds to Eq. (3) and gives each data point the same weight.

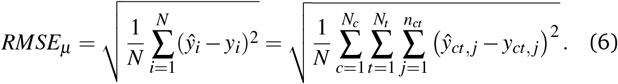

#### Macro-averaged root mean square error (MRMSE)

This one gives each chemical and taxon the same weight:

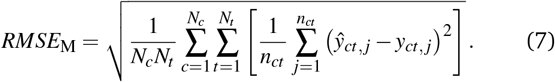

#### Chemical macro-averaged root mean square error (CRMSE)

This one gives each chemical the same weight:

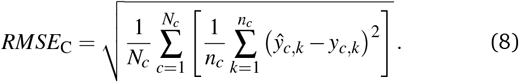

#### Taxon macro-averaged root mean square error (TRMSE)

This one gives each taxon the same weight:

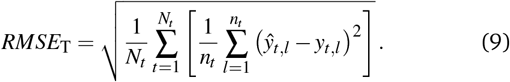

## C Comparison with previous studies – continued

< this section has been moved from the result and discussion section >

Singh *et al*. (2014) implemented ensemble learning models for across-species and across-taxa predictions of log molar effective concentration 50 (EC50). The models were trained on single-species algae data (*P. subcapitata*) using a random train-test-split and 10-fold cross-validation. Since the dataset does not contain repeated experiments, random splitting is adequate. Their decision tree boost and decision tree forest models achieved test RMSEs of 0.56 and 0.64, respectively. They were then used to predict on other algae, crustaceans, fish, and bacteria species ^14^, achieving RMSEs in the range of 0.43 to 0.71. The performance on the fish (medaka, *O. latipes*, 505 data points) is in the same range (0.61 and 0.59, respectively) as the test RMSEs on the algae species. Their better model performance can be attributed to less diverse datasets, *i*.*e*., single-species datasets, less than 800 chemicals in total, and datasets with 40 to 547 chemicals, of which many are shared between data sets. It is unclear how limits, *e*.*g*., LC50 and EC50 larger than a certain value, were processed. The models were based on eight chemical features of which XLogP (logP calculated by an atomic method) and SP-1 (chi simple path descriptor of order 1) were found to be most important.

Toma *et al*. (2021) compiled a dataset on acute and chronic toxicity data for four fish species, algae (*Raphidocelis subcapitata*), and crustaceans (*D. magna*) using data from the Japanese Ministry of Environment and ECOTOX ^12^. For repeated experiments, the geometric mean was calculated and the molar response variables were Box-Cox transformed. Notably, data points with high variability (*±*3 SD from the mean of Box–Cox transformed response values) were excluded. The single-species data subset on acute fish toxicity amounted to 331 chemicals tested on *O. latipes*, for which a RF model, trained after 80:20 train-test-split stratified by LC50 value using 10-fold cross-validation, achieved a test RMSE of 0.87. This is not directly comparable to our work since a different transformation of the response variable was used.

Song *et al*. (2022) used eight single-species datasets, of which five were fish species, to train artificial neural nets to predict the log molar concentration ^13^. The neural nets were trained using 5-fold cross-validation on a training set. The model performance was evaluated on a held-out test set of 20 randomly selected chemicals, leading to *R*^2^ values of 0.54 to 0.72 for the fish data subsets.

## Footnotes

* https://renkulab.io/projects/mltox/mltox-model

