## Supplementary Material for "Machine learning-based prediction of fish acute mortality: Implementation, interpretation, and regulatory relevance"

Journal Name

ARTICLE TYPE

Cite this: DOI: 00.0000/xxxxxxxxxx

Received Date
Accepted Date

DOI: 00.0000/xxxxxxxxxx

1 Data

2 Methods

2.1 Models

3 Results and Discussion

3.1 Data quality & variability

3.2 Modeling results

3.2.1 Validation results

3.2.2 Performance on test set

3.3 Including phylogenetic distances

3.4 Explainability

3.4.1 Residuals

3.4.2 Feature importance

3.4.3 Species sensitivity

References

[a] *Swiss Data Science Center (SDSC), Andreasstrasse 5, Zürich, Switzerland.*
[b] *Eawag, Swiss Federal Institute of Aquatic Science and Technology, Dübendorf, Switzerland.*
[c] *ETH Zürich: Department of Computer Science, Zürich, Switzerland.*
[d] *ETH Zürich: Department of Environmental Systems Science, Zürich, Switzerland.*
[e] *EPF Lausanne, School of Architecture, Civil and Environmental Engineering, Lausanne, Switzerland.*
‡ These authors contributed equally to this work.

Table 1 Response and feature variables. The column name corresponds to the naming in the input datasets.

| Column name | Description | Type of variable |
|---|---|---|
| Response variables | | |
| *result_conc1_mean* | Lethal mass concentration value (in $mg/L$) | continuous |
| *result_conc1_mean_mol* | Lethal molar concentration value (in $mol/L$) | continuous |
| *result_conc1_mean_log* | Lethal mass concentration value after a log10 transformation | continuous |
| *result_conc1_mean_mol_log* | Lethal molar concentration value after a log10 transformation | continuous |
| Experimental features | | |
| *result_obs_duration_mean* | Observation duration (in hours) | ordinal |
| *test_media_type* | Media type | nominal |
| *test_exposure_type* | Exposure type | nominal |
| *result_conc1_type* | Exposure concentration type | nominal |
| Chemical properties | | |
| *chem_mw* | Molecular weight (in $g/mol$) | continuous |
| *chem_ws* | Water solubility (in $mg/L$) | continuous |
| *chem_mp* | Melting point (in $°C$) | continuous |
| *chem_rdkit_clogp* | Octanol-water partition coefficient | continuous |
| Molecular representations | | |
| *chem_MACCS_fp* | Collapsed MACCS fingerprint | binary bits |
| *chem_pcp_fp* | Collapsed PubChem fingerprint | binary bits |
| *chem_Morgan_fp* | Collapsed Morgan fingerprint | binary bits |
| *chem_ToxPrint_fp* | Collapsed ToxPrint fingerprint | binary bits |
| *chem_mol2vec[000-299]* | 300-dimensional mol2vec embedding | continuous |
| *chem_mordred_x* | Mordred features | continuous or ordinal |
| Taxonomic properties | | |
| *tax_eco_climate* | Ecology, climate zone | nominal |
| *tax_eco_ecozone* | Ecology, ecozone | nominal |
| *tax_eco_food* | Ecology, food class | nominal |
| *tax_eco_migrate2* | Ecology, migratory behavior (binary encoding) | binary |
| *tax_lh_amd* | Life history, life span (in $d$) | continuous |
| *tax_lh_licm* | Life history, ultimate body length (in $cm$) | continuous |
| *tax_ps_ampv* | Pseudo-data, energy conductance (in $cm/d$) | continuous |
| *tax_ps_ampkap* | Pseudo-data, allocation fraction to soma | continuous |
| *tax_ps_amppm* | Pseudo-data, volume-specific somatic maintenance cost (in $J/d \cdot cm^3$) | continuous |
| Phylogenetic distance | | |
| stored in a separate file | Phylogenetic distance | continuous |

Table 2 Hyperparameters for the four models. *For the Gaussian process regression hyperparameter, no default values is given in the implementation.

| Model | Hyperparameter | Default | Gridsearch values |
|---|---|---|---|
| LASSO | alpha | 1.0 | [np.round(i, 5) for i in np.logspace(-5, 0, num=26)] |
| RF | n_estimators | 100 | 50, 100, 150, 300 |
| | max_depth | None | 50, 100, 200 |
| | min_samples_split | 2 | 2, 5, 10 |
| | max_samples | None | 0.25, 0.5, 1.0 |
| | max_features | 1.0 | 'sqrt', 1 |
| XGBoost | n_estimators | 100 | 50, 100 |
| | eta | 0.3 | 0.1, 0.2, 0.3 |
| | gamma | 0 | 0, 1, 10 |
| | max_depth | | 3, 6, 9, 12 |
| | min_child_weight | 1 | 1, 3, 5 |
| | subsample | 1 | 0.5, 1. |
| GP | n_inducing | * | 100, 250, 500, 1000 |

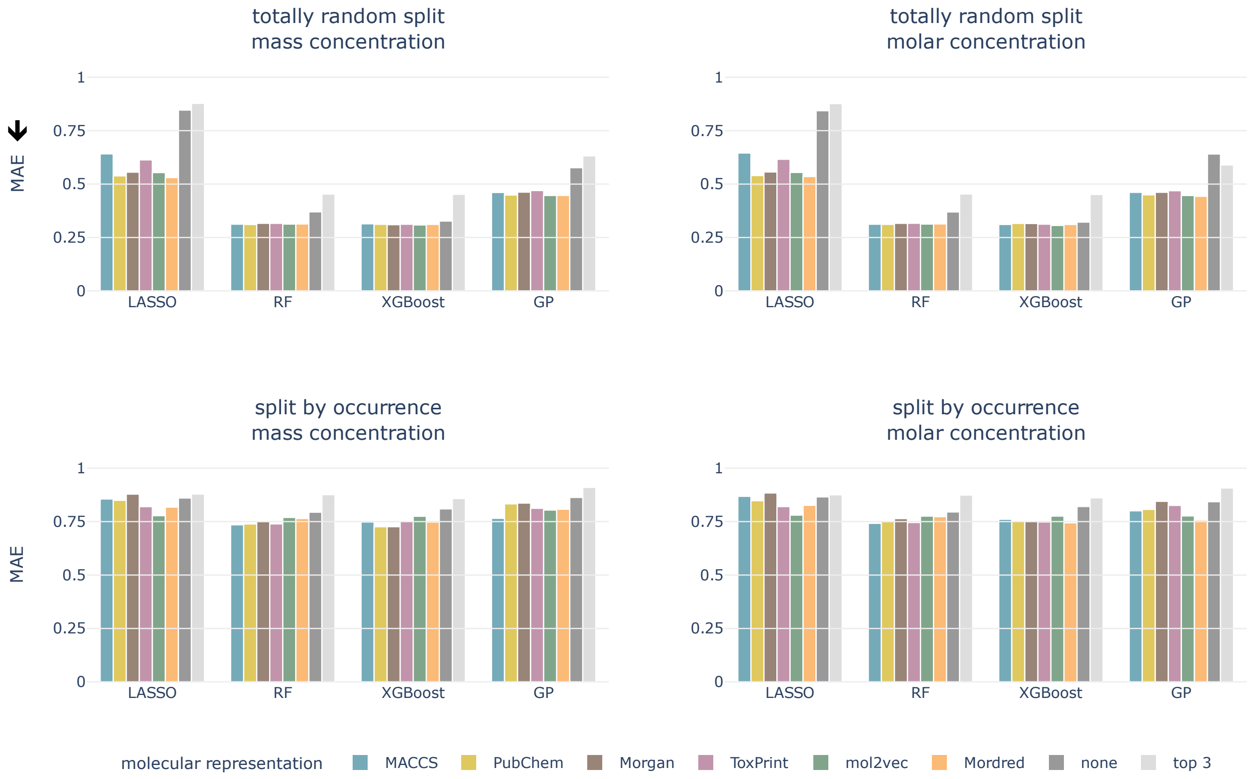


Fig. 1 Cross-validated MAE for both data splittings, concentration types, all models, and molecular representations. Arrow indicates the lower the better.

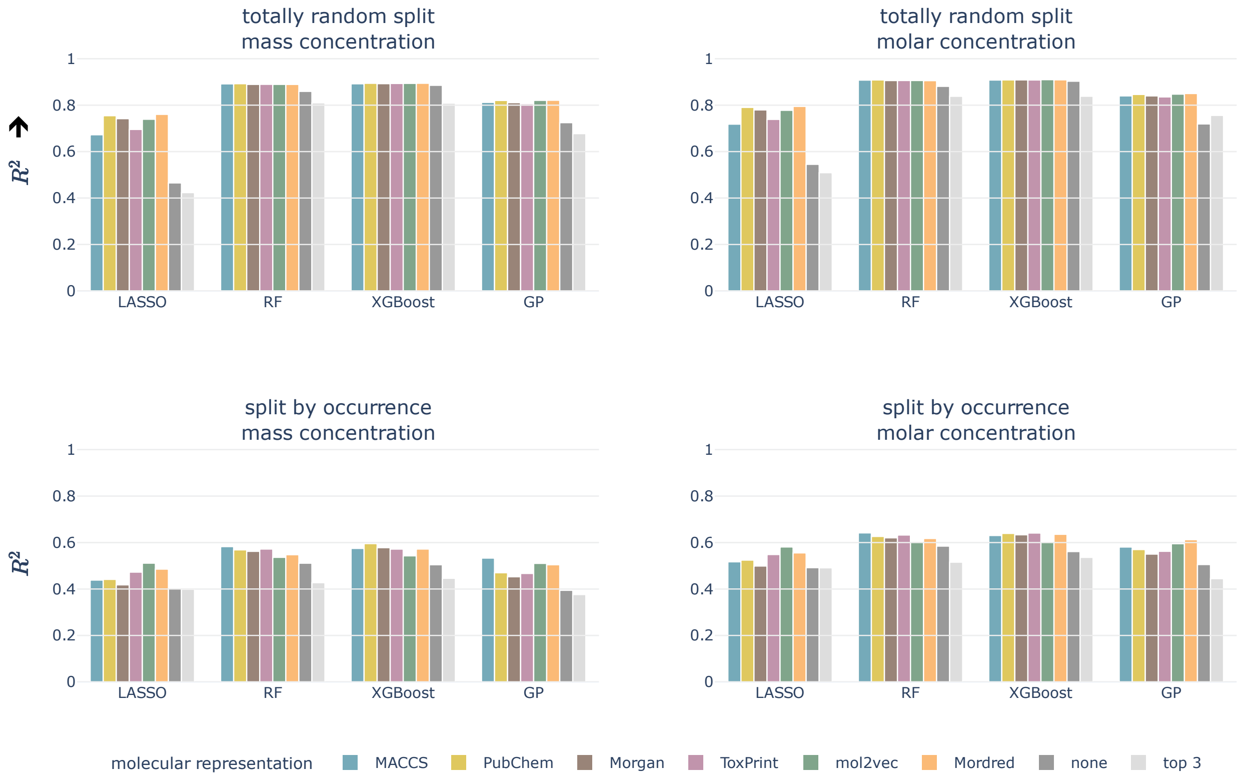


Fig. 2 Cross-validated $R^2$ for both data splittings, concentration types, all models, and molecular representations. Arrow indicates the higher the better.

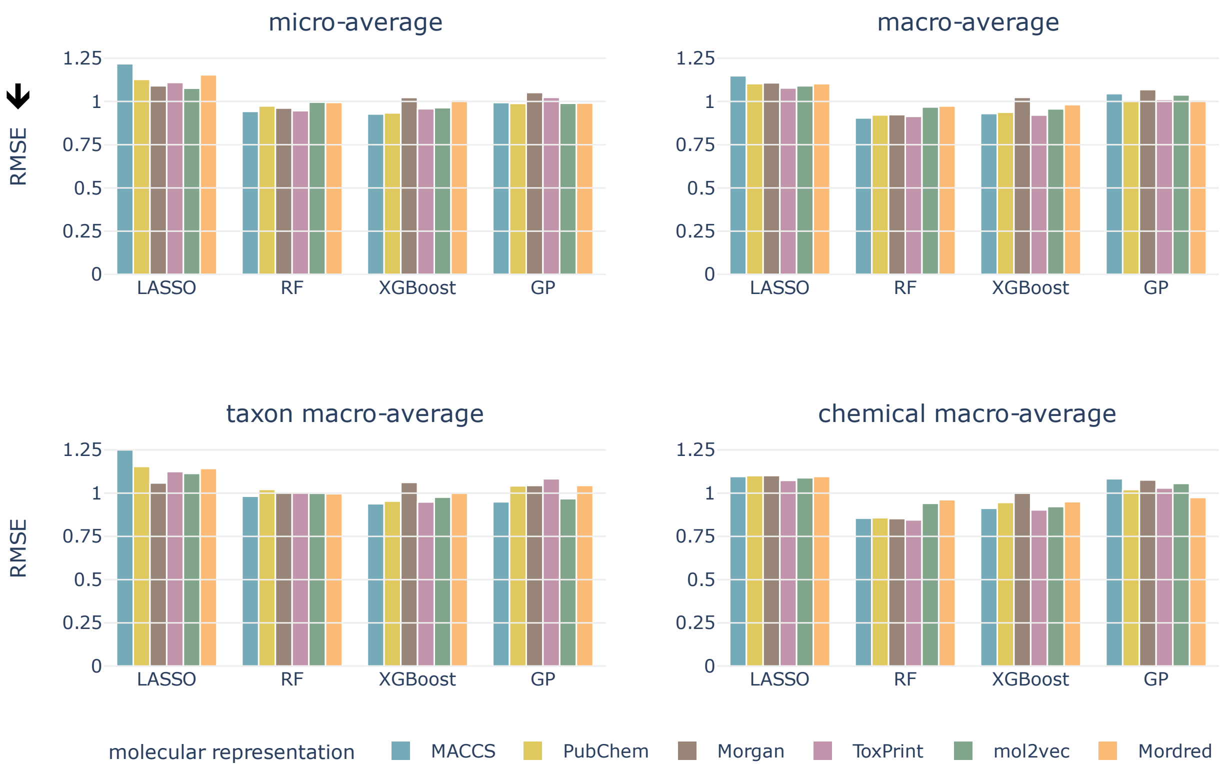


Fig. 3 Test micro- and macro-averaged RMSE for molar LC50, split by occurrence of chemical compounds, all models, and molecular representations. Arrow indicates the lower the better.

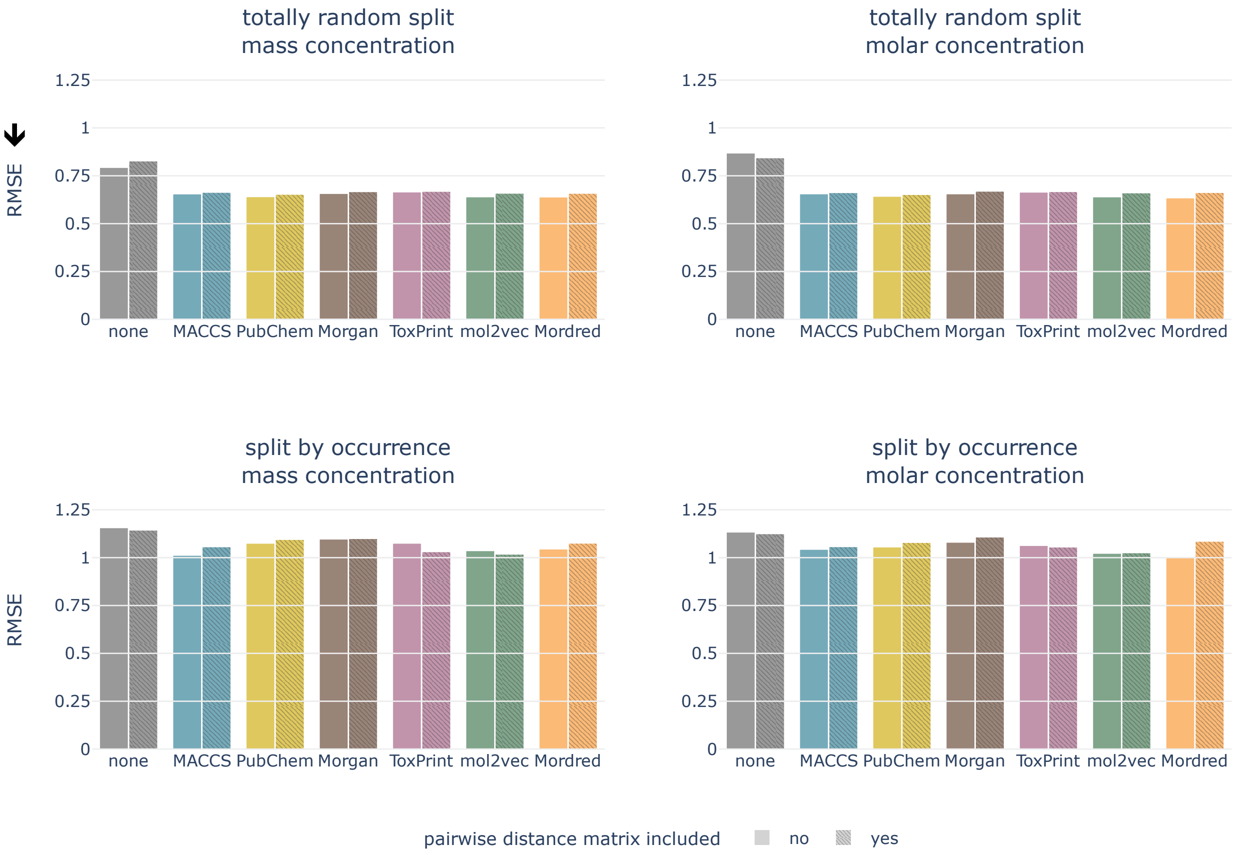


Fig. 4 Including phylogenetic distances: cross-validated RMSE for Gaussian process models, both data splittings, concentration types, and all molecular representations. Arrow indicates the lower the better.

Table 3 Best hyperparameters for LASSO.

| concentration | data split | mol. repr. | alpha |
|---|---|---|---|
| molar | totally random | MACCS | 0.00001 |
| molar | totally random | PubChem | 0.00004 |
| molar | totally random | Morgan | 0.00001 |
| molar | totally random | ToxPrint | 0.00002 |
| molar | totally random | mol2vec | 0.00025 |
| molar | totally random | Mordred | 0.00016 |
| molar | occurrence | MACCS | 0.01585 |
| molar | occurrence | PubChem | 0.02512 |
| molar | occurrence | Morgan | 0.01000 |
| molar | occurrence | ToxPrint | 0.00631 |
| molar | occurrence | mol2vec | 0.01585 |
| molar | occurrence | Mordred | 0.10000 |
| mass | totally random | MACCS | 0.00001 |
| mass | totally random | PubChem | 0.00004 |
| mass | totally random | Morgan | 0.00010 |
| mass | totally random | ToxPrint | 0.00002 |
| mass | totally random | mol2vec | 0.00010 |
| mass | totally random | Mordred | 0.00010 |
| mass | occurrence | MACCS | 0.02512 |
| mass | occurrence | PubChem | 0.02512 |
| mass | occurrence | Morgan | 0.01000 |
| mass | occurrence | ToxPrint | 0.00631 |
| mass | occurrence | mol2vec | 0.01000 |
| mass | occurrence | Mordred | 0.10000 |

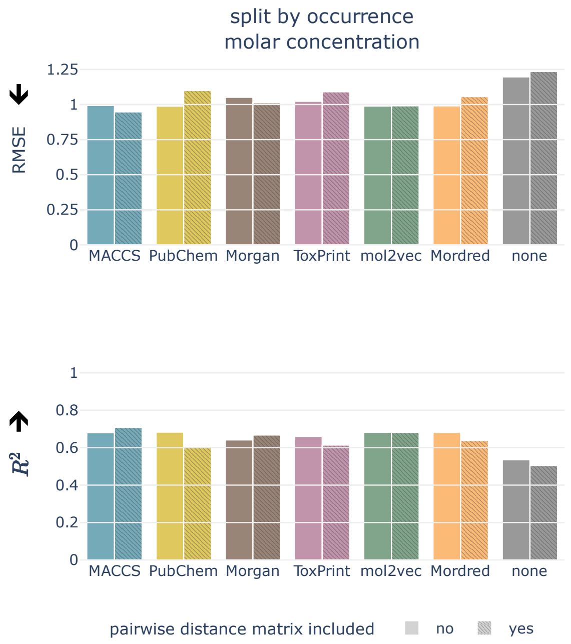


Fig. 5 Including phylogenetic distances: test RMSE and $R^2$ for Gaussian process models, molar LC50, split by occurrence of chemical compounds, and all molecular representations. Arrows indicate the lower/higher the better.

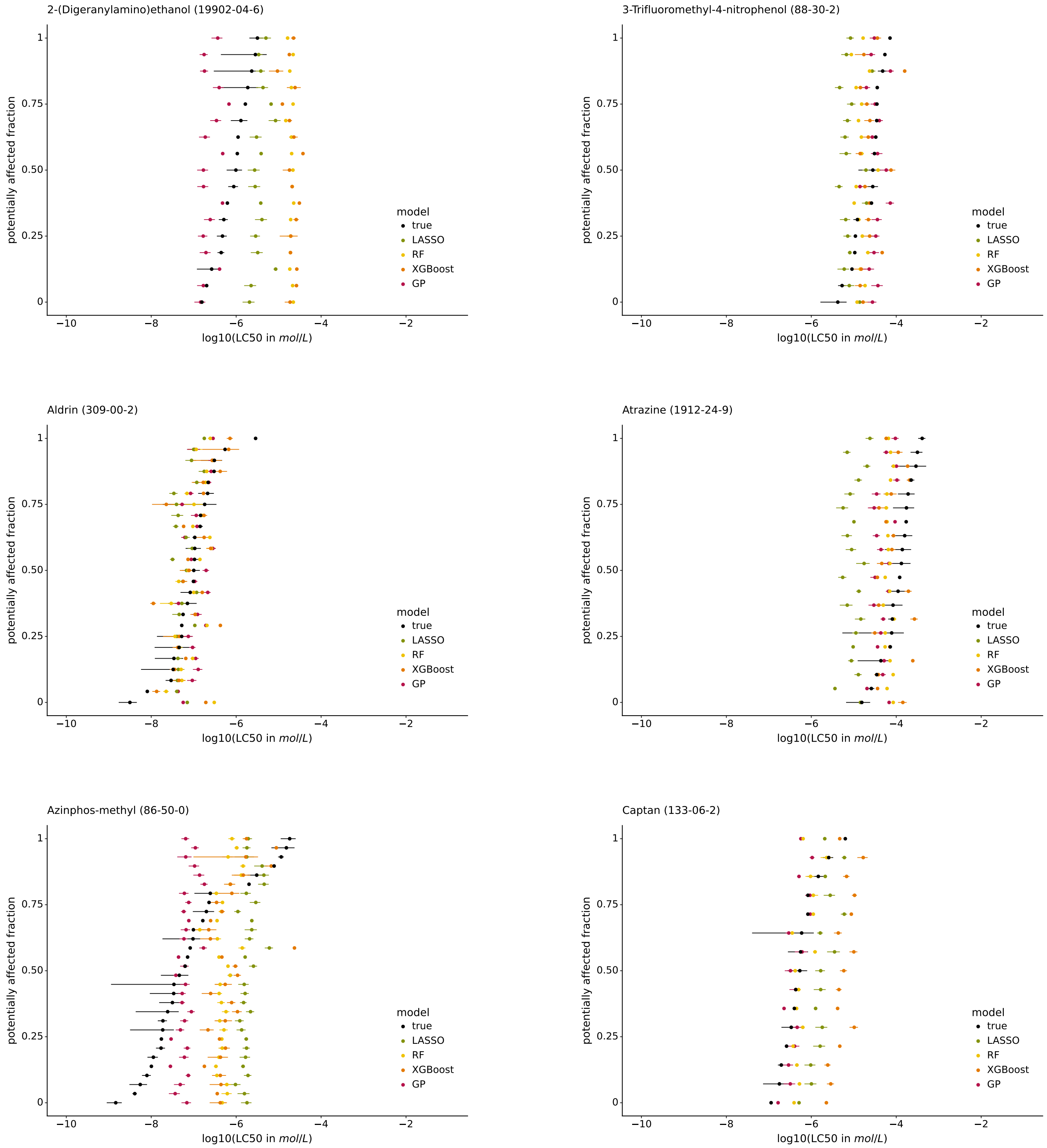


Fig. 6 Species sensitivity distributions (SSDs) of 2-(Digeranylamino)ethanol, 3-Trifluoromethyl-4-nitrophenol, Aldrin, Atrazine, Azinphos-methyl, and Captan.

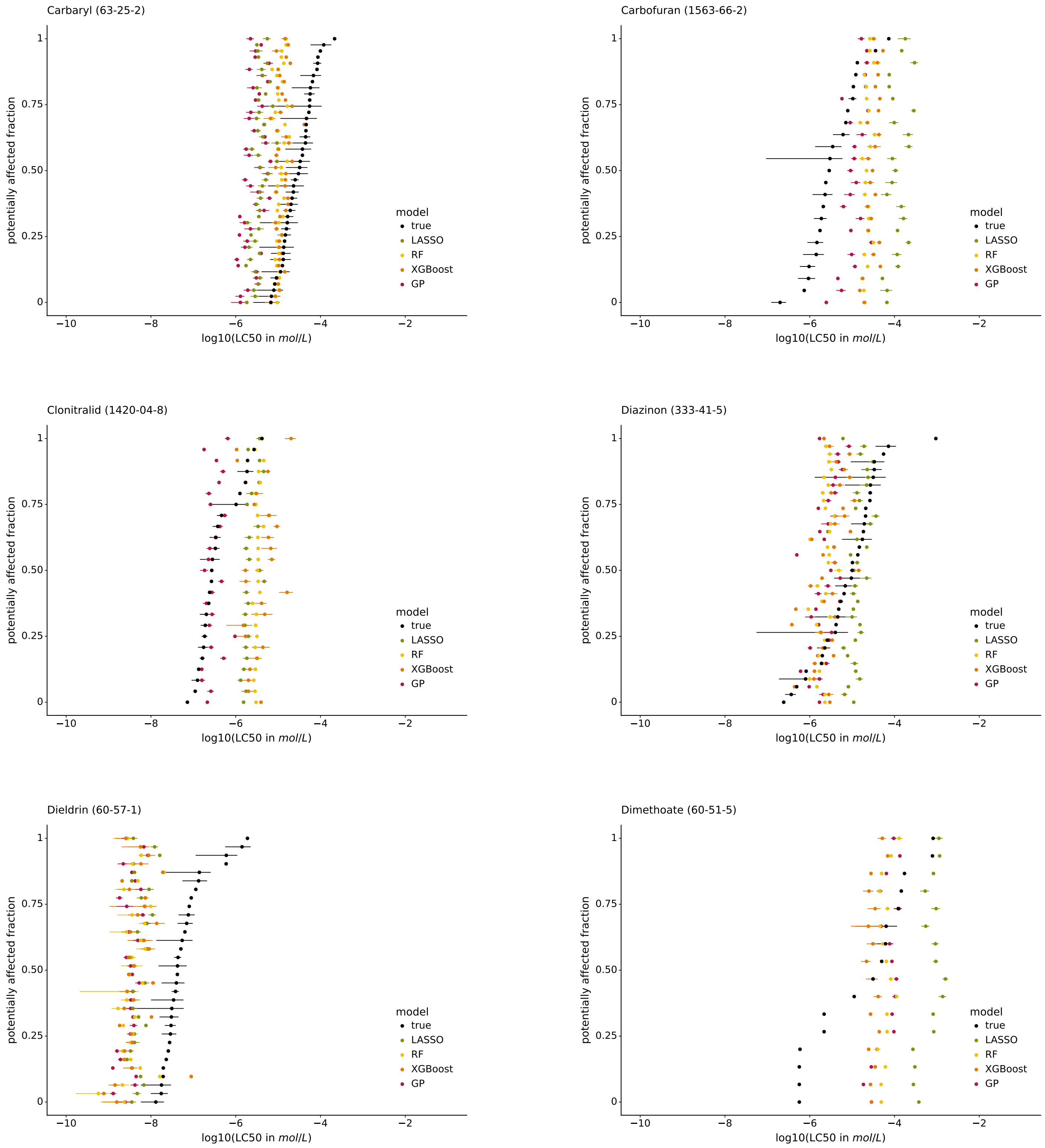


Fig. 7 Species sensitivity distributions (SSDs) of Carbaryl, Carbofuran, Clonitralid, Diazonin, Dieldrin, and Dimethoate.

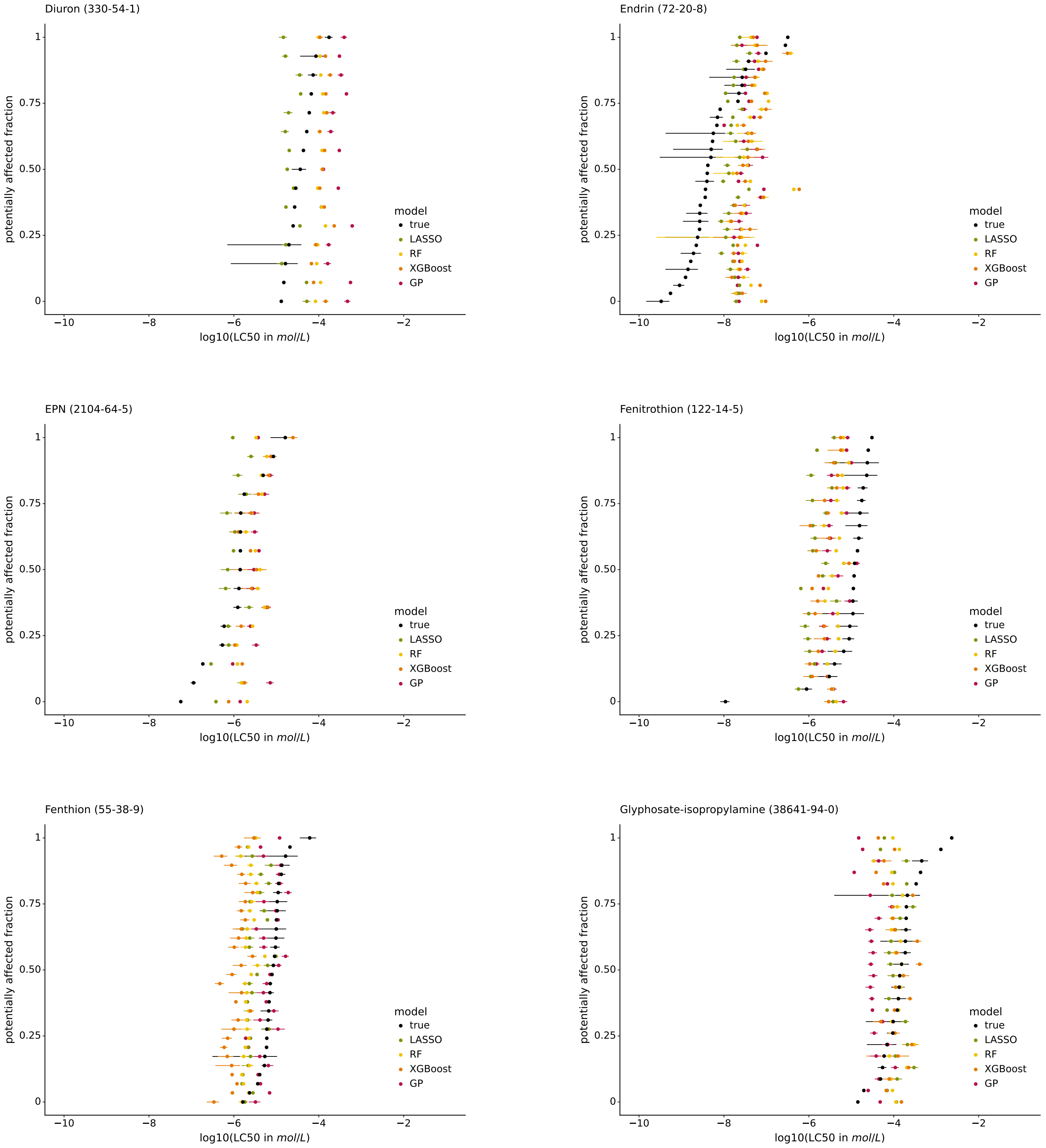


Fig. 8 Species sensitivity distributions (SSDs) of Diuron, Endrin, EPN (Ethyl p-nitrophenyl), Fenitrothion, Fenthion, and Glyphosate-isopropylamine.

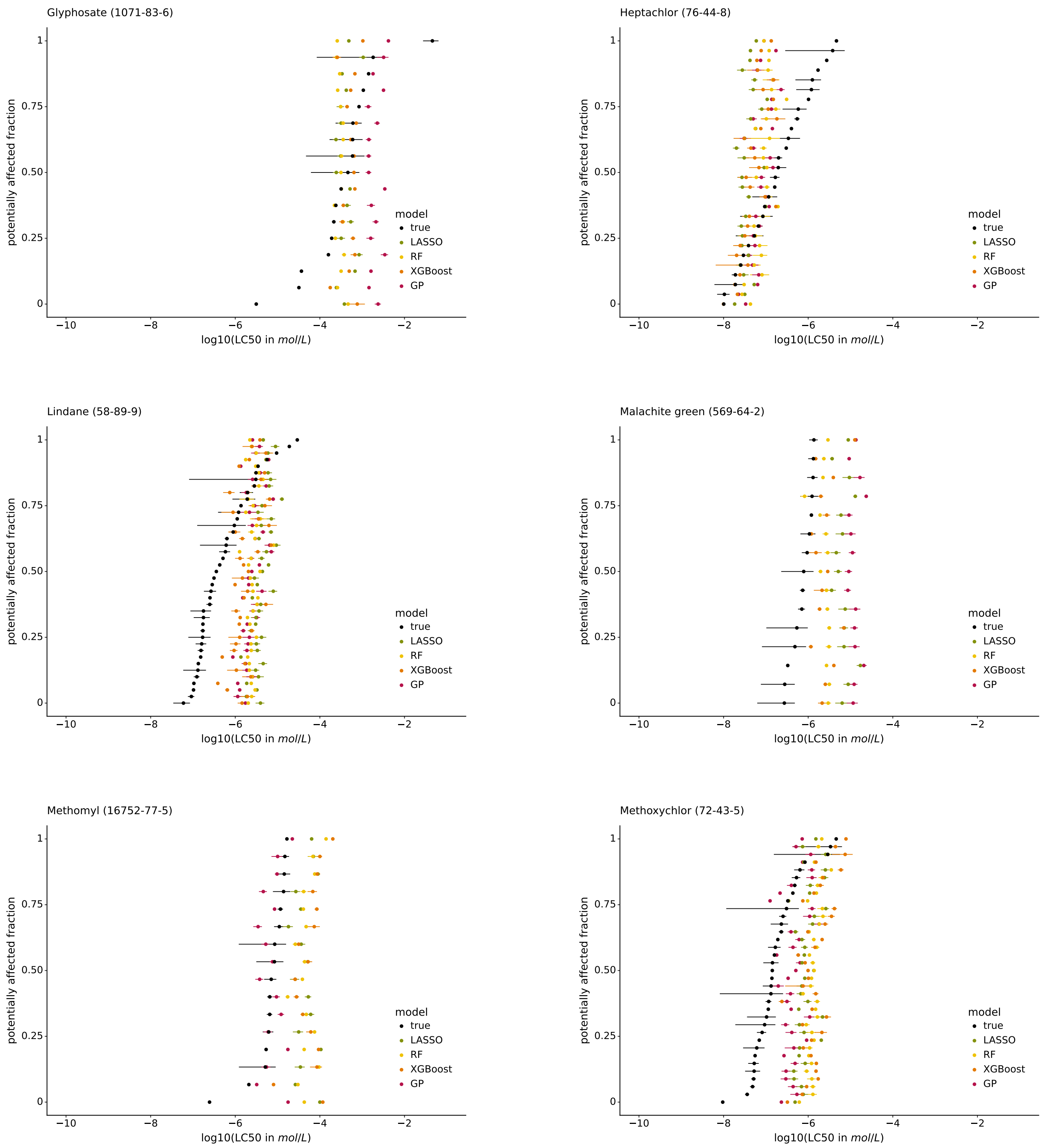


Fig. 9 Species sensitivity distributions (SSDs) of Glyphosate, Heptachlor, Lindane, Malachite green, Methomyl, and Methoxychlor.

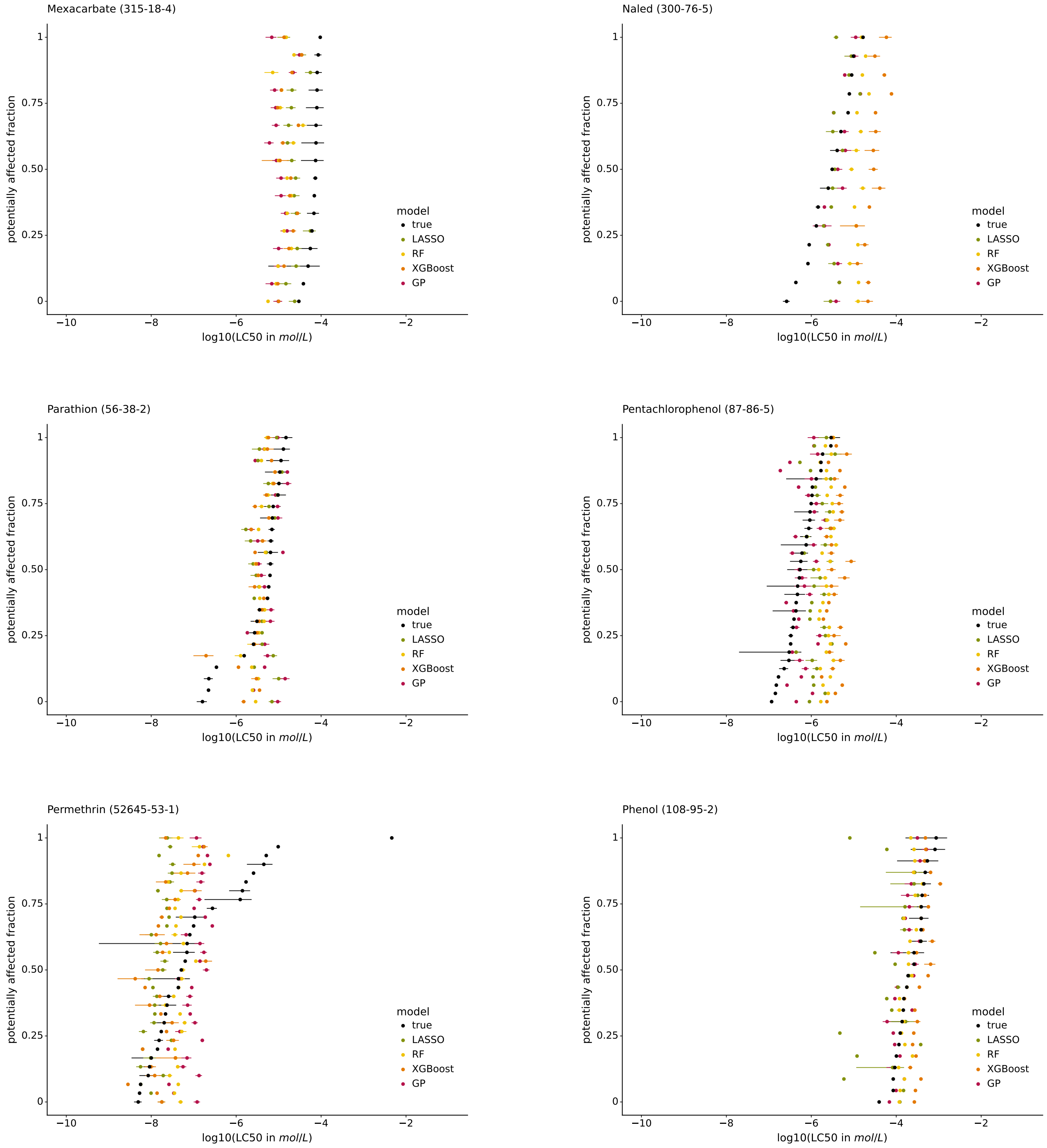


Fig. 10 Species sensitivity distributions (SSDs) of Mexacarbate, Naled, Parathion, Pentachlorophenol, Permethrin, and Phenol.

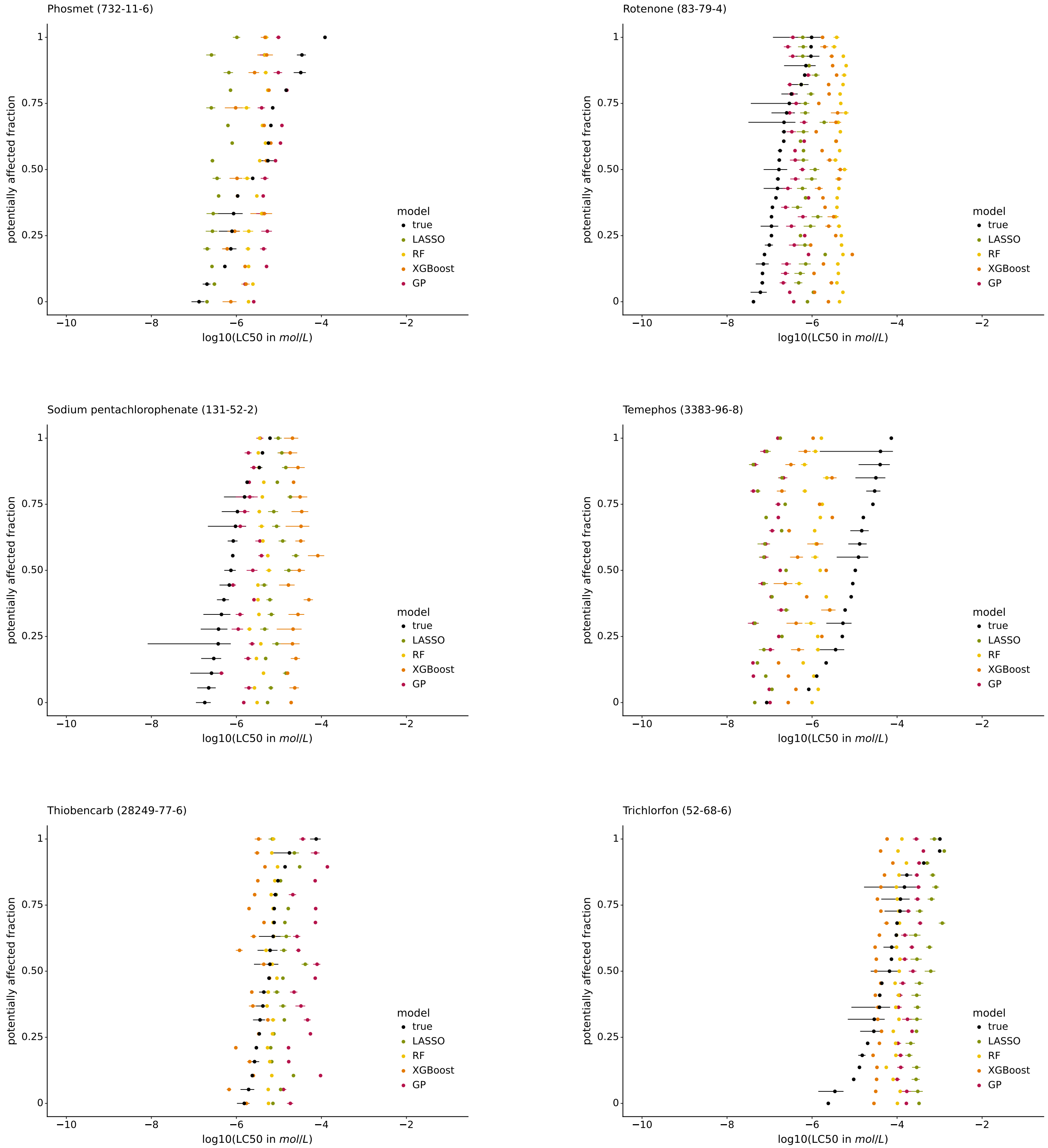


Fig. 11 Species sensitivity distributions (SSDs) of Phosmet, Rotenone, Sodium pentachlorophenate, Temephos, Thiobencarb, and Trichlorfon.

Table 4 Best hyperparameters for random forest. The hyperparameter min_samples_leaf was fixed to the default value of 1.

| concentration | data split | mol. repr. | n_estimators | max_depth | max_samples | min_samples_split | max_features |
|---|---|---|---|---|---|---|---|
| molar | totally random | MACCS | 300 | 50 | 1.0 | 2 | sqrt |
| molar | totally random | PubChem | 150 | 200 | 1.0 | 2 | sqrt |
| molar | totally random | Morgan | 300 | 50 | 1.0 | 2 | sqrt |
| molar | totally random | ToxPrint | 300 | 100 | 1.0 | 2 | sqrt |
| molar | totally random | mol2vec | 300 | 200 | 1.0 | 2 | sqrt |
| molar | totally random | Mordred | 300 | 200 | 1.0 | 2 | sqrt |
| molar | occurrence | MACCS | 300 | 200 | 1.0 | 2 | sqrt |
| molar | occurrence | PubChem | 150 | 50 | 1.0 | 2 | sqrt |
| molar | occurrence | Morgan | 300 | 100 | 1.0 | 2 | sqrt |
| molar | occurrence | ToxPrint | 50 | 200 | 1.0 | 2 | sqrt |
| molar | occurrence | mol2vec | 300 | 50 | 1.0 | 2 | sqrt |
| molar | occurrence | Mordred | 50 | 50 | 0.25 | 2 | sqrt |
| mass | totally random | MACCS | 300 | 100 | 1.0 | 2 | sqrt |
| mass | totally random | PubChem | 300 | 50 | 1.0 | 2 | sqrt |
| mass | totally random | Morgan | 300 | 200 | 1.0 | 2 | sqrt |
| mass | totally random | ToxPrint | 300 | 50 | 1.0 | 2 | sqrt |
| mass | totally random | mol2vec | 300 | 200 | 1.0 | 2 | sqrt |
| mass | totally random | Mordred | 300 | 100 | 1.0 | 2 | sqrt |
| mass | occurrence | MACCS | 50 | 200 | 1.0 | 2 | sqrt |
| mass | occurrence | PubChem | 100 | 200 | 0.5 | 2 | sqrt |
| mass | occurrence | Morgan | 150 | 50 | 1.0 | 2 | sqrt |
| mass | occurrence | ToxPrint | 100 | 50 | 1.0 | 2 | sqrt |
| mass | occurrence | mol2vec | 50 | 200 | 1.0 | 2 | sqrt |
| mass | occurrence | Mordred | 150 | 200 | 1.0 | 2 | sqrt |

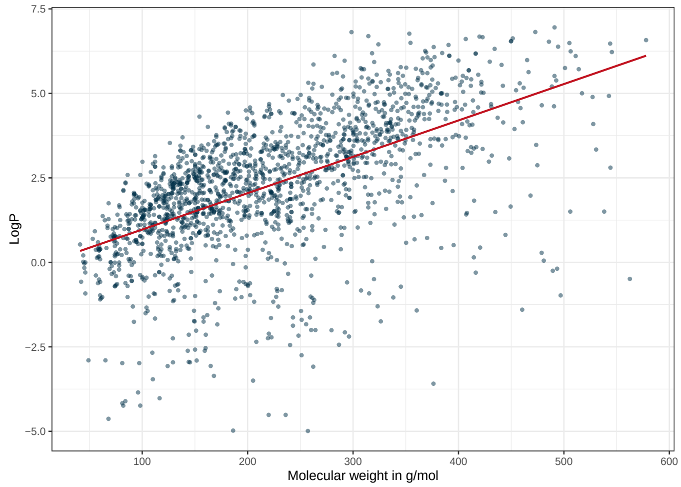


Fig. 12 Correlation of molecular weight and logP for the chemicals in the ADORE "t-F2F" challenge. The red line indicates a linear regression model fit to the data points.

Table 5 Best hyperparameter for XGBoost.

| concentration | data split | mol. repr. | n_estimators | eta | gamma | max_depth | min_child_weight | subsample |
|---|---|---|---|---|---|---|---|---|
| molar | totally random | MACCS | 100 | 0.100000 | 0 | 12 | 1 | 1.0 |
| molar | totally random | PubChem | 100 | 0.100000 | 0 | 12 | 3 | 1.0 |
| molar | totally random | Morgan | 100 | 0.200000 | 0 | 12 | 5 | 1.0 |
| molar | totally random | ToxPrint | 100 | 0.200000 | 0 | 12 | 3 | 1.0 |
| molar | totally random | mol2vec | 100 | 0.100000 | 0 | 12 | 3 | 1.0 |
| molar | totally random | Mordred | 100 | 0.100000 | 0 | 12 | 3 | 1.0 |
| molar | occurrence | MACCS | 100 | 0.100000 | 1 | 9 | 5 | 0.5 |
| molar | occurrence | PubChem | 100 | 0.300000 | 0 | 3 | 3 | 1.0 |
| molar | occurrence | Morgan | 100 | 0.300000 | 1 | 3 | 3 | 1.0 |
| molar | occurrence | ToxPrint | 100 | 0.200000 | 0 | 6 | 5 | 1.0 |
| molar | occurrence | mol2vec | 100 | 0.100000 | 1 | 6 | 3 | 1.0 |
| molar | occurrence | Mordred | 100 | 0.100000 | 0 | 6 | 1 | 0.5 |
| mass | totally random | MACCS | 100 | 0.100000 | 0 | 12 | 1 | 0.5 |
| mass | totally random | PubChem | 100 | 0.300000 | 0 | 9 | 1 | 1.0 |
| mass | totally random | Morgan | 100 | 0.200000 | 0 | 12 | 3 | 1.0 |
| mass | totally random | ToxPrint | 100 | 0.200000 | 0 | 12 | 5 | 1.0 |
| mass | totally random | mol2vec | 100 | 0.200000 | 0 | 9 | 1 | 1.0 |
| mass | totally random | Mordred | 100 | 0.300000 | 0 | 9 | 5 | 1.0 |
| mass | occurrence | MACCS | 100 | 0.200000 | 0 | 6 | 3 | 1.0 |
| mass | occurrence | PubChem | 100 | 0.300000 | 0 | 3 | 5 | 1.0 |
| mass | occurrence | Morgan | 100 | 0.300000 | 1 | 3 | 3 | 0.5 |
| mass | occurrence | ToxPrint | 100 | 0.200000 | 10 | 3 | 5 | 0.5 |
| mass | occurrence | mol2vec | 100 | 0.100000 | 1 | 6 | 3 | 0.5 |
| mass | occurrence | Mordred | 100 | 0.100000 | 1 | 6 | 5 | 1.0 |

Table 6 Best hyperparameters for GP regression.

| concentration | data split | mol. repr. | n_inducing |
|---|---|---|---|
| molar | totally random | MACCS | 1000 |
| molar | totally random | PubChem | 1000 |
| molar | totally random | Morgan | 1000 |
| molar | totally random | ToxPrint | 1000 |
| molar | totally random | mol2vec | 1000 |
| molar | totally random | Mordred | 1000 |
| molar | occurrence | MACCS | 250 |
| molar | occurrence | PubChem | 500 |
| molar | occurrence | Morgan | 100 |
| molar | occurrence | ToxPrint | 500 |
| molar | occurrence | mol2vec | 250 |
| molar | occurrence | Mordred | 1000 |
| mass | totally random | MACCS | 1000 |
| mass | totally random | PubChem | 1000 |
| mass | totally random | Morgan | 1000 |
| mass | totally random | ToxPrint | 1000 |
| mass | totally random | mol2vec | 1000 |
| mass | totally random | Mordred | 1000 |
| mass | occurrence | MACCS | 250 |
| mass | occurrence | PubChem | 250 |
| mass | occurrence | Morgan | 100 |
| mass | occurrence | ToxPrint | 500 |
| mass | occurrence | mol2vec | 100 |
| mass | occurrence | Mordred | 1000 |
