## Supplementary Material – Self-assessment for "Machine learning-based prediction of fish acute mortality: Implementation, interpretation, and regulatory relevance"

**REFORMS Checklist for reporting ML-based science**

### Module 1: Study goals

1a. Population or distribution about which the scientific claim is made.

- The chemicals and species, which are available in dataset
    - For chemicals, we provide basic properties, molecular representations, the chemical ontology (ClassyFire) and functional use information. The dataset contains mainly organic molecules.
    - For species, we provide information from the Add my Pet database as well as phylogenetic distances.
    - See Schür et al. (2023) for details.

1b. Motivation for choosing this population or distribution (1a.).

- data availability, governed by performed tests, relevance of fish for ecotoxicology and high motivation to establish an alternative for fish-based acute toxicity testing.
- mol2vec filtering

1c. Motivation for the use of ML methods in the study.

- to capture non-linear relationships
- to integrate various (types of) features
- to show the potential and limitations of ML

### Module 2: Computational reproducibility

2a. Dataset used for training and evaluating the model along with link or DOI to uniquely identify the dataset.

- Modeling is based on the ADORE dataset
    - Schür, Christoph, Lilian Gasser, Fernando Perez-Cruz, Kristin Schirmer, und Marco Baity-Jesi. „A Benchmark Dataset for Machine Learning in Ecotoxicology“. *Scientific Data* 10, Nr. 1 (18. Oktober 2023): 718. https://doi.org/10.1038/s41597-023-02612-2).
    - The paper proposes modeling challenges and delivers the necessary datasets and train-test-splits. Here, we focus on the t-F2F challenge (i.e., making predictions on 140 fish species).
    - The dataset is deposited and freely available through an institutional repository: https://doi.org/10.25678/0008c9

2b. Code used to train and evaluate the model and produce the results reported in the paper along with link or DOI to uniquely identify the version of the code used.

- The code used to train and evaluate the models can be found here: https://renkulab.io/gitlab/mltox/mltox-model

2c. Description of the computing infrastructure used.

- Hardware infrastructure: CPU, GPU, RAM, disk space etc.
    - VM: 8 CPUs, 1 GPU, 43GB RAM, 24GB disk space
    - Laptop: 8 CPUs, 0 GPU, 15GB RAM, 8GB disk space
- Operating system.
    - VM: Ubuntu 20.04
    - Laptop: Ubuntu 20.04
- Software environment: Programming language and version, documentation of all packages used along with versions and dependencies (e.g., through a requirements.txt file).
    - Python 3.11
    - See environment.yml in mltox-model repository
- An estimate of the time taken to generate the results.
    - 2-3 weeks

2d. README file which contains instructions for generating the results using the provided dataset and code.

- The modeling repository contains a README with a detailed description how to set it up and run the necessary scripts.

2e. Reproduction script to produce all results reported in the paper[1].

- We do not provide a reproduction script. To produce our results, the following needs to be done (relevant scripts start with a two-digit number):
    - Hyperparameter tuning and cross-validation per model: scripts 11, 12, 13, 14
    - Evaluate cross-validation per model: scripts 21, 22, 23, 24
    - Generate test results per model: scripts 31, 32, 33, 34
    - Evaluate all results: script 46
    - Evaluate residuals: script 47

- Evaluate macro-averages: script 48
- Evaluate feature importances and generate SSDs: script 53-54

**Module 3: Data quality**

3a. Source(s) of data, separately for the training and evaluation datasets (if applicable), along with the time when the dataset(s) are collected, the source and process of ground-truth annotations, and other data documentation.

- The base dataset is compiled and filtered from ECOTOX database (September 2022 Version) to capture acute mortality in fish, crustaceans and algae under laboratory conditions. It mainly contains experimental data.
- It is extended with taxonomic (phylogenetic distances and species-related properties) and chemical data (chemical properties and molecular representations).
- It is provided with training, test and cross-validation splits (within the training data).
- Curation process is described in the dataset paper: https://doi.org/10.1038/s41597-023-02612-2

3b. Distribution or set from which the dataset is sampled (i.e., the sampling frame).

- Not applicable. We compiled the dataset from experimental data available in ECOTOX.

3c. Justification for why the dataset is useful for the modeling task at hand.

- To our knowledge, this is the largest compilation of relevant ecotoxicity data to predict acute mortality in fish, crustaceans and algae.

3d. The definition of the outcome variable of the model along with descriptive statistics, if applicable.

*(The outcome variable is also known as the dependent variable, the target variable, the output variable or the predicted variable).*

- log10 transformed effective concentration 50 (EC50), either mass (in mg/L) or molar (mol/L)

3e. Number of samples in the dataset.

- In the t-F2F challenge: 26,114 samples

3f. Percentage of missing data, split by class for a categorical outcome variable.

- The features used for modeling do not have missing values.

3g. Justification for why the distribution or set from which the dataset is drawn (3b.) is representative of the one about which the scientific claim is being made (1a.).

- We compiled the available data on acute mortality in three relevant taxonomic groups and filtered it to contain samples representing standard experimental conditions.

**Module 4: Data preprocessing**

4a. Identification of whether any samples are excluded with a rationale for why they are excluded.

- The reasoning for the data pre-processing is described extensively in Schür et al. (2023).
- Given the large number of samples in the dataset, we did not check every entry. We ask users of the datasets to report errors through an issue on the gitlab repository of ADORE.

4b. How impossible or corrupt samples are dealt with.

- Corrupted samples were either corrected or deleted. See methods section of Schür et al. (2023) for details.

4c. All transformations of the dataset from its raw form (3a.) to the form used in the model, for instance, treatment of missing data and normalization.

- See methods section of Schür et al. (2023) and of this paper.

**Module 5: Modeling**

5a. Detailed descriptions of all models trained, including:

- All features used in the model (including any feature selection).
  - see section 2.4 and supplemental Table 1
- Types of models implemented (e.g., Random Forests, Neural Networks).
  - LASSO, Random Forest, XGBoost, Gaussian Process regression
- Loss function used.
  - quadratic loss

5b. Justification for the choice of model types implemented.

- Baseline model: LASSO
- State-of-the-art in ecotoxicology: Random Forest, XGBoost
- Our contribution: Gaussian Process regression

5c. Method for evaluating the model(s) reported in the paper, including details of train-test splits or cross-validation folds.

- The data is split according to occurrence of chemical compounds (see Schür et al. (2023) and this paper for details)

5d. Method for selecting the model(s) reported in the paper.

- We selected the models with the best performance in hyperparameter tuning via gridsearch. On the training data, 5-fold cross-validation is performed and the RMSE across the five folds is averaged. The model with the lowest averaged RMSE (from the five folds) is selected and evaluated on the test set.

5e. For the model(s) reported in the paper, specify details about the hyperparameter tuning:

- Range of hyper-parameters used and a justification for why this range is reasonable.
  - see supplemental Table 2
  - justification: standard ranges
- Method to select the best hyper-parameter configuration.

- Grid-search

● Specification of all hyper-parameters used to generate results reported in the paper.

- see supplemental Table 2

5f. Justification that model comparisons are against appropriate baselines.

- Since it is a new dataset, we compare several models with each other. The linear LASSO model can be considered a baseline model.

**Module 6: Data leakage**

6a. Justification that pre-processing (Section 4) and modeling (Section 5) steps only use information from the training dataset (and not the test dataset).

- Potential sources are discussed and critically evaluated in Schür et al. (2023).
- Please refer to the data leakage checklist.

6b. Methods to address dependencies or duplicates between the training and test datasets (e.g. different samples from the same patients are kept in the same dataset partition).

- With the occurrence split, the same chemical only occurs in one of the cross-validation folds or in the test data.
- Since the dataset is also highly skewed with respect to species, we could not perform a split by chemical and species. Therefore, the same species is likely to occur in several folds and/or the test set.

6c. Justification that each feature or input used in the model is legitimate for the task at hand and does not lead to leakage.

- We only use general chemical, experimental and taxonomic properties. We have excluded features that could be considered to risk data leakage such as modes of action, chemical ontology, and functional use data.

**Module 7: Metrics and uncertainty**

7a. All metrics used to assess and compare model performance (e.g., accuracy, AUROC etc.). Justify that the metric used to select the final model is suitable for the task.

- RMSE, MAE and R2. They are standard metrics for regression.

7b. Uncertainty estimates (e.g., confidence intervals, standard deviations), and details of how these are calculated.

- No uncertainty estimates were used.

7c. Justification for the choice of statistical tests (if used) and a check for the assumptions of the statistical test.

- No statistical tests were used.

**Module 8: Generalizability and limitations**

8a. Evidence of external validity.

- Since we are not happy with the results obtained on the internal dataset, we did not perform external validation.

8b. Contexts in which the authors do not expect the study's findings to hold.

- Outside the tested toxicity range.
- For chemicals outside the model's applicability domain.
- For non fish species and for fish species, that are very different from the species present in the dataset.

---

[1] Note that this is a high bar for computational reproducibility. It might not be possible to provide such a script—for instance, if the analysis is run on an academic computing cluster, or if the dataset does not allow for programmatic download.

| | | |
|---|---|---|
| **Kapoor et al. 2023: Leakage and reproducibility** | | |
| **12 pitfalls** | | |
| [L1] Lack of clean separation of training and test data | | We provide data splits for all challenges. For all splits but the “totally random” split, chemicals either appear in the training or test set. Species appear in both sets, since a separation by both is not feasible as the dataset is highly skewed with regard to chemicals and species. |
| [L1.1] No test set | | We have a separate test set for each challenge. |
| [L1.2] Pre-proc. on train-test | | We do not perform imputation or over/undersampling. Standardscaling is done on the training set. |
| [L1.3] Feature selection on training and test set | | We perform feature selection on the training set. |
| [L1.4] Duplicates in datasets | | There are no duplicates in the dataset. Nevertheless, the same species appear both in the training and test set (see L1). |
| [L2] Model uses features that are not legitimate | | Features either describe the chemical properties or structure, the taxonomy or experimental conditions. |
| [L3] Test set is not drawn from the distribution of scientific interest | | For the “occurrence” split, training and test sets are from the same distribution, which is of intereset since we want to do interpolation. It is unclear, how well our model performs for yet unseen chemicals. |
| [L3.1] Temporal leakage | N/A | Not applicable. |
| [L3.2] Nonindependence between train and test samples | | For the “occurrence” split, chemicals are either in the training or test set. But the same species can be found both in the training and test set (see L1). |
| [L3.3] Sampling bias in test distribution | | Difficult to judge. |
| **Other issues identified in our survey** | | |
| Computational reproducibility issues | | We share code, dataset, as well as the dependencies needed to rerun the code. |
| Data quality issues | | We generated the dataset to the best of our knowledge. Nevertheless, future studies could find our dataset to be flawed. We are transparent about obvious limitations of both our dataset and the models. |
| Metric choice issues | | We report the standard metrics for regression (R2, RMSE, MAE). |
| Use of standard datasets | N/A | We make our dataset available as a benchmark for others. |

| | | |
|---|---|---|
| | Legend | |
| | | Fulfilled |
| | | Partly fulfilled |
| | | Not fulfilled |
| | N/A | Not applicable |

| **Cronin et al. 2019: Identification and description of the uncertainty, variability and bias and influence in QSARs for toxicity prediction** | | | Legend: Low (blue); Moderate (orange); High (red); N/A: Not applicable |
|---|---|---|---|
| ID | Area of Uncertainty, Variability, Bias or Influence | Assignment of Uncertainty, Variability, Bias or Influence | Reasoning |
| **Model Creation** | | | |
| *1.1 Definition of Chemical Structures* | | | |
| 1.1a | Accuracy of chemical structure | Moderate | Structures well-defined but not accounting for any isomerism. Multiple substance identifiers provided. |
| 1.1b | Assessment of significant impurities or mixtures | Low | Impurities and mixtures were removed in the generation of the ADORE dataset. |
| 1.2 Biological Data | | | |
| 1.2a | Quality of individual studies in the data set | High | We removed experimental properties related to non-standard designs in the preprocessing (e.g., only durations of 24, 48, 72, and 96 h), which increases likelihood of results coming from standardized test guidelines. Nonetheless, we cannot conclusively determine how many data points come from similar experimental designs. However, data has been evaluated before inclusion |

| | | | |
|---|---|---|---|
| | | | into the ECOTOX described in (Olker et al. (2022)) and Add my Pet (process described at https://www.bio.vu.nl/thb/deb/deblab/add_my_pet/) databases. |
| 1.2b | Consistency of the data set including comparability of data | | Varied test guidelines and sources of data, but measures were undertaking to reduce variability and to increase conformity with standardized test guidelines. |
| 1.2c | Checking of toxicological data | | We manually checked some of the entries but not at random but rather different kinds of outliers (extreme outcome or other variables and outliers withing their respective distribution of similar experiments). |
| 1.2d | Error associated with biological data | | In the paper characterizing the curation process and the dataset itself (Schür et al. (2023)) and in the given study, we describe the biological variability through the range of outcomes associated with experiments where species, chemical, and experimental variables overlap. |
| 1.2e | (if required) Units of concentration known, stated and appropriate for use | | Harmonized to appropriate units, provide both mass- and mole-based concentrations. |
| 1.2f | (If appropriate) Nominal or measured concentrations | | Concentrations are not always measured, often nominal concentrations are used; inconsistent across the dataset. |
| 1.2g | Internal exposure known | | Either nominal or measured concentration of the medium are recorded, internal concentrations of the organisms throughout the experiments are not included. |
| *1.3 Measurement and / or Estimation of Physico-Chemical Properties and Structural Descriptors* | | | |
| 1.3a | Measurement of physico-chemical properties | N/A | No measured properties used, only predicted. |
| 1.3b | Calculation of properties and 2-D descriptors | | Well-characterised software providing unambiguous properties. Where calculation was unclear it is stated transparently (i.e., pKa) |
| 1.3c | Calculation of 3-D descriptors | N/A | No 3-D descriptors used. |

| | | | |
|---|---|---|---|
| 1.3d | Software utilised | | Full details of software and code provided. |
| 1.3e | Definition of molecular fragments | | For many of the molecular representations the individual bits are documented. Links to the definitions are given in the dataset paper. |
| *1.4 Creation of the Data Set for QSAR Modelling* | | | |
| 1.4a | Data set is complete | | Yes, data points with missing values are removed. |
| 1.4b | Data set has appropriate variation in potency (quantitative) or balance of actives vs inactives (qualitative) | | Good variation in potency (e.g. several log units). |
| 1.4c | Selection of training set data | | Training set selected without bias (i.e., split by occurrence of chemical). |
| 1.4d | Training set homogeneity | | Density plot for important chemical properties are presented to display the homogeneity of training and testing data. |
| 1.4e | Suitable training and test sets defined and utilised | | Appropriate training and test splits. |
| *1.5 Modelling Approach* | | | |
| 1.5a | How appropriate is the modelling approach for the endpoint and to deal with the complexity / non-linearity of the data | | Regression analysis is an appropriate modelling approach for the endpoint |
| **Description of the QSAR Model** | | | |

| | | | |
|---|---|---|---|
| *2.1 Description of Model* | | | |
| 2.1a | Documentation and reporting | | Model fully defined |
| 2.1b | Data set is complete and described | | All data are provided and described in the ADORE paper. |
| 2.1c | Transparency of the model | | Model is transparent in terms of the algorithm. |
| *2.2 Statistical Performance* | | | |
| 2.2a | Statement of statistical fit, performance and predictivity | | Full description of model performance (CV and test errors) |
| 2.2b | Interpretation of statistical fit etc with respect to biological error (see Criterion 1.2d) | | Statistical performance is significant but not overfitted |
| *2.3 Applicability Domains* | | | |
| 2.3a | Chemical applicability domain of model | | Fully defined in terms of relevant physico-chemical properties and structure |
| 2.3b | Mechanistic applicability domain of model | N/A | The selected endpoint LC50 is unspecific and not limited to a certain mode of action. We discuss this as a limitation and how it impacts the reliability of the model prediction. |
| 2.3c | Biological applicability domain of model | | We limit our dataset and model to the taxonomic group of fish (140 species). We evaluate the cross-species extrapolation ability of our model across this group. Metabolism-related data is included through the Add my Pet parameters. |

| | | | |
|---|---|---|---|
| *2.4 Mechanistic Relevance, Interpretability and Transparency* | | | |
| 2.4a | Mechanistic justification | | The selected endpoint LC50 is very unspecific. Hence, many different mechanisms are present. We generated an overview of the predicted modes of action present using several available QSAR models (Verhaar, Ecosar, Oasis) and the majority of chemicals could not be classified according to their mode of action. We discuss this in the dataset paper as well as in the present study. |
| 2.4b | Presence / availability of other and supporting information | | Functional use and chemical ontology data is included with the dataset that can link modeling outcomes with potential mechanisms of action. Users can use the given identifiers to produce their own mode of action predictions/classifications with freely available QSAR models (e.g. included in the QSAR toolbox). However, we do not endorse the use of such information as modeling features because, in our oppinion, that would constitute a kind of data leakage. |
| 2.4c | Relevance of descriptors to mechanism of action / AOP | | See 2.4a and 2.4b. The inclusion of feature importance analysis could provide an intuition of relevant chemical properties and substructures. |
| *2.5 Adequate coverage of ADME effects* | | | |
| 2.5a | Metabolism and / or effect of significant metabolites have been considered | | Currently there is not large-scale quantitative data for biotransformation products and their toxicity available, hence this is not included in our model. The Add my Pet parameters provide a linkage to inter-species variability in organism metabolism, but do not establish a link related to biotransformation. |

| | | | |
|---|---|---|---|
| 2.5b | Toxicokinetics have been addressed in the model | | Currently there is not large-scale quantitative data for biotransformation products and their toxicity available, hence this is not included in our model. |
| **Application of the QSAR Model** | | | |
| *3.1 Documentation and Reproducibility* | | | |
| 3.1a | Reproducibility of the models | | Model transparently and fully documented |
| 3.1b | Reproducibility of the prediction | | We document the distribution of model residuals across highly-represented chemicals and species and discuss this in the paper. |
| *3.2 Usability* | | | |
| 3.2a | Implementation of the model | | Fully implemented into software |
| 3.2b | Software accessibility | | Software is publicly and freely available |
| 3.2c | Software transparency | | Software algorithm is transparent |
| 3.2d | Relative cost | | Standardized guideline tests with vertebrates require 42-350 animals per test at an annual cost upwards of $39 mio. (for fish and birds combined) (Mittal et al. 2022). Hence, a model would be more cost-effective by several orders of magnitude. |
| 3.2e | Sustainability | | The model is made available through an institutional repository, but long-term support is hindered through limited staff continuity. |
| 3.2f | Maintenance and support | | The model is made available through an institutional repository, but long-term support is hindered through limited staff continuity. |
| 3.2g | Intellectual Property | N/A | Since the models are not sufficiently good, no licence is needed. |

| | | | |
|---|---|---|---|
| 3.2h | Ownership | | Ownership and contact information provided. |
| 3.2i | Ethics | | No ethical concerns |
| *3.3 Relevance* | | | |
| 3.3a | Heterogeneity and density of chemical space | | Well populated and distributed chemical space. We acknowledge a selection bias in the available toxicity data. Additionally, the use of mol2vec limits the chemical space to a certain degree. |
| 3.3b | Relevance of the predicted endpoint for the regulatory risk assessment purpose/protection goal | | Fit for stated purpose. Likely to provide an estimate that could support hazard identification. We discuss the relevance for the regulatory context. |
| 3.3c | Adequacy | | Adequate for stated purpose. Likely to provide an estimate that could support e.g. hazard identification. We discuss the potential relevance and the limitations for the regulatory context in the paper. |
| 3.3d | Extrapolation and relevance to humans | N/A | Our work is focused on ecotoxicology (i.e. fish). Hence, human relevance is limited, but also not the scope of this paper. |
| 3.3.e | Extrapolation and relevance to environmental biota | | Relevant to environmental biota. Across-taxa-extrapolation needs to be further explored, but is also not the scope of this paper. |
